# Restructuring of an asymmetric neural circuit during associative learning

**DOI:** 10.1101/2023.01.12.523604

**Authors:** Leo T.H. Tang, Garrett A. Lee, Steven J. Cook, Jacquelin Ho, Cassandra C. Potter, Hannes E. Bülow

**Author notes:** These authors contributed equally to this work (Leo T.H. Tang, Garrett A. Lee).

## Abstract

Asymmetric brain function is common across the animal kingdom and involved in language processing, and likely in learning and memory. What regulates asymmetric brain function remains elusive. Here, we show that the nematode *Caenorhabditis elegans* restructures an asymmetric salt sensing neural circuit during associative learning. Worms memorize and prefer the salt concentration at which they were raised in the presence of food through a left-biased network architecture. When conditioned at elevated salt concentrations, animals change the left-biased to a right-biased network, which explains the changed salt-seeking behavior. The changes in circuit architecture require new synapse formation induced through asymmetric, paracrine insulin-signaling. Therefore, experience-dependent changes in asymmetric network architecture rely on paracrine insulin signaling and are fundamental to learning and behavior.

## Introduction

Lateralized brain function in humans was first recognized by Pierre Paul Broca in 1865 [1]. His and subsequent studies showed that language and speech processing are generally localized to the left frontal and temporal cortical lobes, establishing that lateralized brain function is a common feature of the human brain [2, 3]. Asymmetries in brain function, however, are not limited to language and speech; several other processes are associated with lateralized brain functionality, including emotion and perception [4, 5]. Moreover, defects in lateralized brain function have been implicated with neuropsychiatric conditions such as autism spectrum disorders and schizophrenia, among others [6-8].

Lesions in the left lateral cortex due to strokes, tumors or traumatic brain injury result in language and speech deficits, but these faculties can often be relearned through the recruitment of both ipsilateral, perilesional and contralateral homologous brain regions [9, 10]. These findings suggest that asymmetric neural circuits are not static but can be plastic. This is also supported by functional magnetic resonance imaging (fMRI) studies demonstrating the recruitment of asymmetric connectivity of brain circuits during learning of a motor task [11]. Lateralized nervous system function and structure are not limited to humans and common to most animals, yet its functional significance or the underlying mechanisms that establish and mediate lateralization are not well understood [2, 12]. Here we use the nematode *Caenorhabditis elegans*, which displays neuronal asymmetries in sensory neurons on the anatomical as well as functional level [13, 14] and for which the full synaptic connectome has been established [15, 16], to uncover molecular pathways involved in the plasticity of nervous system lateralization during associative learning.

## Results and Discussion

### Synaptic output of ASE salt-sensing neurons correlates with associative salt-learning

*C. elegans* is capable of associative learning. Worms recall and seek the temperature or the salt concentration at which they were raised in the presence of food when placed on a temperature or salt gradient [17, 18]. Consistent with these studies, we found that *C. elegans* animals grown on agar plates with 33 mM sodium chloride (NaCl) in the presence of food (hereafter termed naïve), preferred agar with 33 mM NaCl over agar with 100 mM NaCl in a quadrant-choice assay (Fig. 1A-B). After conditioning for 12 hours on 100 mM NaCl in the presence of food (conditioned), worms preferred 100 mM in the choice assay (Fig. 1A-B). Thus, within 12 hours worms form an associative memory of and preference for the salt concentration they experienced with food. *C. elegans* senses salt primarily through the pair of gustatory ASE neurons in the head (Fig. 1C), the left/right members of which serve distinct functions and display asymmetric cell fates [13]. While the left-side member, ASEL, primarily responds to acute increases in cations such as Na^+^, the right-side ASER neuron responds to an acute decrease in anions, such as Cl^-^ [19]. Although the functions of these neurons have been previously studied in attractive and aversive learning paradigms [18, 20], their synaptic connectivity in this context has not been analyzed.

**Fig. 1.**
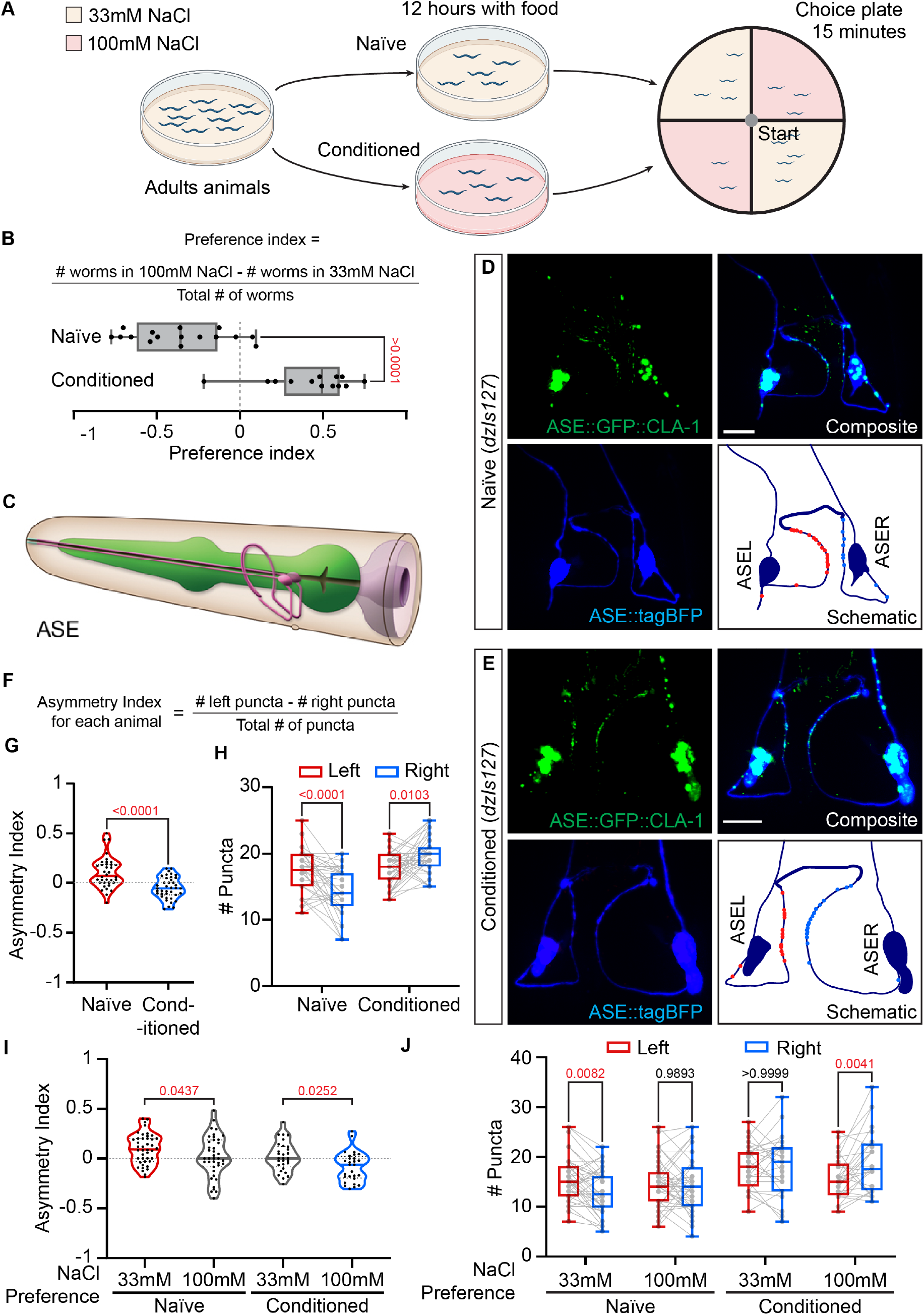
Salt preference correlates with ASE asymmetric synaptic connectivity. **(A)** Schematic of salt conditioning protocol and choice assay **(B)** Formula for calculating preference indices, graphed for naïve (N = 16) and conditioned (N = 13) animals. P value labeled on graph, two-tailed unpaired t test. (**C-D**) Representative Images of animals carrying the *dzIs127* transgene (*Is[flp-6p::tagBFP+flp-6p::GFP::cla-1]*) labeling ASE neurons in blue and ASE presynaptic puncta in green in a naïve **(C)** and a conditioned animal (D). Scale bar = 10 μm. In the schematic, red and blue puncta indicate location of ASE presynaptic puncta on the left and on the right respectively. (**E**) Diagram showing the anatomy of the ASE neurons (www.wormatlas.org). (**F-H**) Formula for calculating the asymmetry indices for each animal. The number of ASE GFP::CLA-1 puncta of naïve (N = 33) and conditioned (N = 39) animals were counted on each side and the asymmetry indices for each animal is calculated. Asymmetry indices are plotted in (G), analyzed with two-tailed unpaired t test. Number of puncta are plotted in (H) was analyzed with two-way matched ANOVA with Sidak’s multiple comparison. (**I**-**J**) Naïve and conditioned animals are segregated by their salt preference after the choice assay shown in (A). Asymmetry indices based on GFP::CLA-1 puncta plotted in (I), and number of puncta in (J). (Naïve: N = 42 for 33 mM preference, N = 39 for 100 mM preference, Conditioned, N = 29 for 33 mM preference, N = 28 for 100 mM preference). P value in (I) and (J) from unmatched two-way ANOVA and two-way matched ANOVA with Sidak’s multiple comparison respectively.

To study synaptic connectivity and asymmetry during associative salt learning, we visualized all presynaptic specializations in ASE neurons by transgenically expressing a cell-specific GFP::CLA-1 active zone marker [21] and counted the number of active zone puncta as a proxy for ASE total synaptic output (Fig. 1D-E). We calculated an asymmetry index for each animal, defined as the ratio of the difference between the number of puncta on the left versus right and the total number of puncta (Fig. 1F). Positive and negative numbers indicate biased connectivity for a given animal on the left or right, respectively. Additionally, we determined the average number of puncta on the left and right side in a population of animals. We found that naïve animals displayed a left bias in ASE connectivity. A majority of animals showed more presynaptic active zones along the processes of the left ASEL neuron compared to the right ASER neuron, although there was variability within the population (Fig. 1D, G-H, ASEL: 17.36 ± 0.54 puncta, ASER:14.28 ± 0.61 puncta). In conditioned animals, the difference was reversed. The right ASER neuron contained more active zones than the left ASEL (Fig. 1E, G-H, ASEL puncta: 17.60 ± 0.41 vs ASER puncta: 19.56 ± 0.36).

Since salt conditioning changed salt preference behavior, we asked whether the behavior of individual animals correlated with the observed change in asymmetric connectivity. We found that naïve animals that had chosen the 33 mM NaCl concentration at which they were raised displayed a left bias in the number of presynaptic active zones (Fig. 1I-J). In contrast, naïve animals that had ‘erroneously’ chosen the 100 mM quadrant showed symmetric numbers of presynaptic densities (Fig. 1I-J). Conversely, animals conditioned for 12 h on 100 mM NaCl that chose the 100 mM quadrant displayed a right bias in ASE connectivity, whereas animals that had ‘erroneously’ chosen the 33 mM quadrant showed again symmetric connectivity (Fig. 1I-J). These findings show that (1) the main salt-sensing neurons ASEL and ASER contain different numbers of presynaptic active zones, (2) an associative salt learning paradigm reverses the left/right bias in connectivity; and (3) lateralization of connectivity is correlated with animal behavior.

### ASE>AWC synaptic connectivity is plastic and dependent on experience

Recent work identified asymmetric synaptic connectivity of ASE salt-sensing neurons [16]. We therefore revisited the postsynaptic connectivity of ASEL and ASER neurons in the connectomes [16], and found that only connectivity between ASE and AWC olfactory neurons displayed left-biased asymmetry in connectivity (Table S1). To test whether the asymmetric connectivity observed by the GFP::CLA-1 active zone marker is reflective of ASE>AWC connectivity, we used iBLINC technology [22], which allows specific labeling of the ASE>AWC synaptic connection [16]. We found that 81% of naïve animals displayed a left-biased ASE>AWC connection, which after conditioning for 12 hours on 100 mM NaCl changed to 65% of the animals displaying a right-biased ASE>AWC connection (Fig. S1A). The ASE>AWC connection in naïve animals contained 5.88 ± 0.52 puncta on the left vs 2.88 ± 0.42 on the right, which changed to right biased connectivity after conditioning with 3.88 ± 0.30 puncta on the left vs 5.37 ± 0.37 on the right (Fig. 2A-D). The difference in synaptic puncta between ASEL and ASER neurons was similar between naïve and conditioned animals using the presynaptic active zone marker GFP::CLA-1 or the iBLINC reporter (3.08 ± 0.74 and 3.00 ± 0.42, respectively, in naïve animals, and −1.97±0.64 and −1.49±0.55, respectively, in conditioned animals), suggesting that ASE>AWC is the predominant asymmetric synaptic connection of ASE neurons. We used the ASE>AWC iBLINC reporter to further characterize the plasticity of this asymmetric connection.

**Fig. 2.**
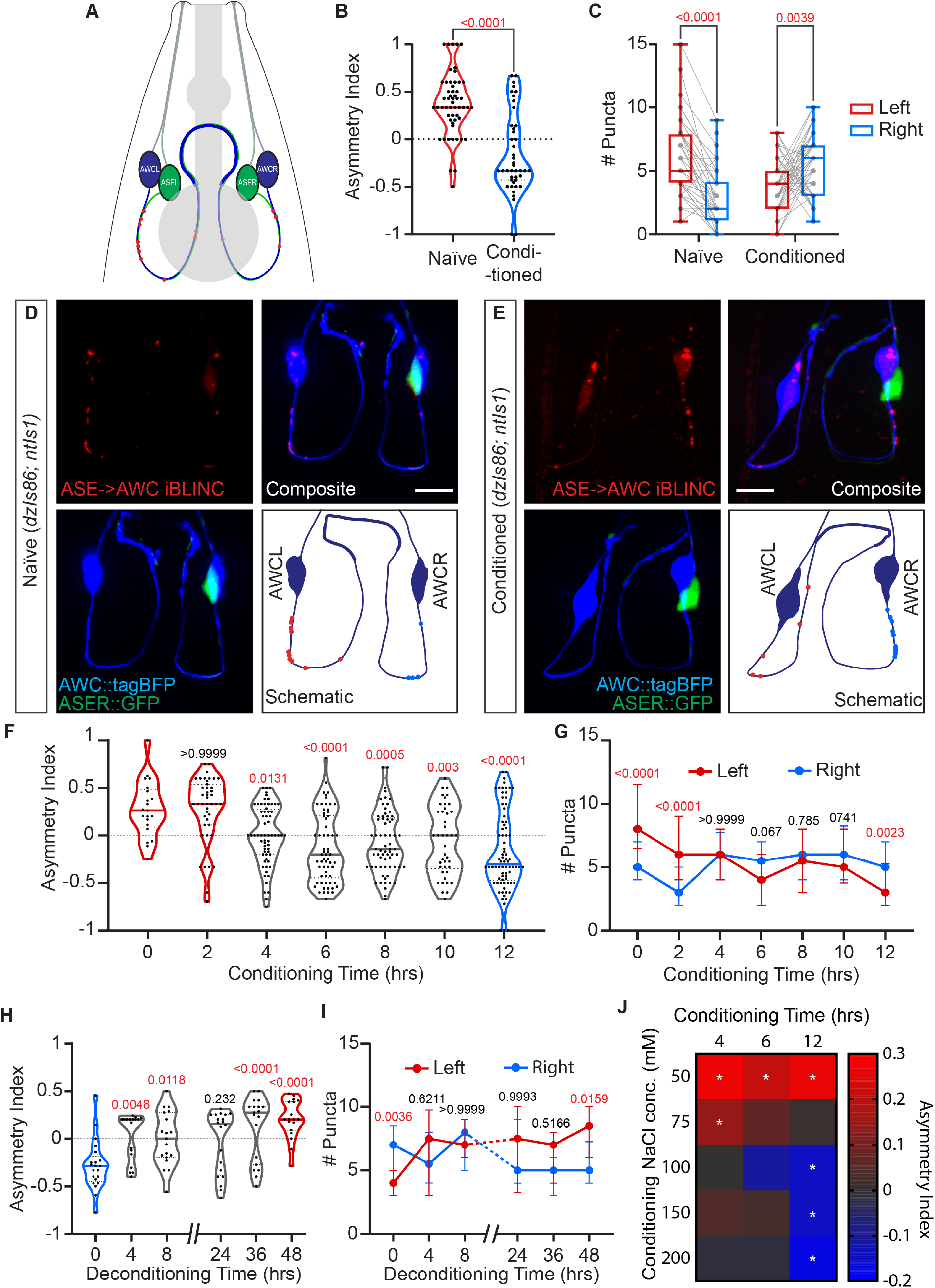
ASE asymmetric connectivity is mediated by experience of changed salt concentration. (**A**) Schematic of a dorsal view of ASE>AWC synaptic connectivity. (**B-C**) Asymmetry indices (B) and number of puncta (C) comparing iBLINC puncta between naïve animals (N = 20) and 100 mM NaCl conditioned animals (N = 43). (B) was analyzed with two-tailed unpaired t test. (C) was analyzed with two-way matched ANOVA with Sidak’s multiple comparison. (**D-E**) Representative images of naïve (D) and conditioned (E) animals carrying transgenes *dzIs86 (ASE>AWC iBLINC + AWC::tagBFP)* and *ntIs1 (ASER::GFP)*. iBLINC puncta are red, AWC neurons are in blue and ASER in green. Scale bar = 10 μm. In the schematic, red and blue puncta indicate location of iBLINC puncta on the left and on the right respectively. (**F-G**) Asymmetry indices (F) and number of puncta (G) showing iBLINC asymmetry at the times indicated during 100 mM NaCl conditioning. N = 21 for 0 hrs, 43 for 2 hrs, 64 for 4 hrs, 62 for 6 hrs, 66 for 8 hrs, 42 for 10 hrs and 77 for 12 hrs. (F) was analyzed with unmatched one-way ANOVA. P value from Tukey multiple comparison test against hour 0 are labled. (G) was analyzed with two-way matched ANOVA with Sidak’s multiple comparison. (**H-I**) Asymmetry indices (G) and number of puncta (H) showing iBLINC at the times indicated during deconditioning (back to 33 mM NaCl after 100 mM conditioning). N = 21 for 0 hrs, 18 for 4 hrs, 19 for 8 hrs, 20 for 24 hrs, 19 for 36 hrs, and 18 for 48 hrs. (H) was analyzed with unmatched one-way ANOVA. P value from Tukey multiple comparison test against hour 0 are indicated. (I) was analyzed with two-way matched ANOVA with Sidak’s multiple comparison. (**J**) Asymmetry indices plotted as heatmap for different time points and concentrations of NaCl conditioning. N > 16 for each square. Analyzed with two-way matched ANOVA for each time point, with Sidak’s multiple comparison.

A time course showed that ASE>AWC connectivity becomes symmetric after 4 hours of conditioning but requires 12 hours until it becomes right-biased (Fig. 2F). This was due to an increase of synapses on the right after 4 hours, and a more gradual decrease of synapses on the left, which suggests that two different mechanisms may contribute to the changes in connectivity (Fig. 2G). The change in connectivity was reversible, recovering to a left-bias after 48 hours when conditioned animals were placed back on 33mM NaCl plates with food (Fig. 2H-I). The generation of new synapses on the right side (but not the decrease in synapses on the left side) was dependent on *de novo* protein synthesis, as blocking translation prevented only the increase of right-side synapses during conditioning (Fig. S1B-C). The change in connectivity was dosage dependent as smaller increases of salt concentration failed to change the asymmetry, or merely symmetrized the connection (Fig. 2J). Moreover, the change in connectivity was dependent on the relative increase of NaCl rather than the absolute concentration of NaCl (Fig. 2J, Fig. S1D-E). Animals raised at 100 mM NaCl displayed a left bias (just like those raised at 33 mM), and their synapses only became symmetric or right-biased when they were conditioned at 150 mM or 200 mM NaCl for 12 hours, respectively (Fig. S1D-E). The change in connectivity was not the result of changes in osmolarity, because exposure of worms to an osmotic increase equivalent to 100 mM NaCl using glycerol had no effect on ASE>AWC connectivity (Fig. S1F-G). Lastly, the observed effects appeared specific to the ASE>AWC connection, as other unrelated synaptic connections of sensory neurons, (e.g. the AFD>AIY sensory to interneuron synaptic connection) were impervious to changes in salt exposure (Fig. S1H). Collectively, these findings show that (1) synaptic connectivity between the ASE salt-sensing and AWC olfactory neurons is plastic, and that (2) synaptic “hardwiring” can change in response to experience.

### ASE>AWC asymmetric connectivity is established during larval development and dependent on presynaptic cell fate

We next asked how this asymmetry in connectivity is established. We found that freshly hatched larvae displayed symmetric connectivity between ASE and AWC neurons, which became asymmetric during larval development and into adulthood. This asymmetry was retained during adulthood for at least four days (Fig. S2A-B). Therefore, ASE>AWC asymmetric connectivity is established through the asymmetrical addition of synapses during larval stages rather than through pruning or during embryonic development.

Both left/right ASE neurons and AWC neurons display asymmetric cell fates due to dedicated transcriptional programs. ASEL and ASER neurons express asymmetric cell fates, e.g. through the asymmetric expression of different receptor-type guanyl cyclases [13]. By contrast, AWC neurons are antisymmetric, i.e. display a random asymmetry, as defined by the mutually exclusive expression of G-protein coupled receptors [14]. We found that ASE>AWC connectivity maintained a left bias if both presynaptic ASEs adopted a developmental ASER fate. In contrast, a right bias was observed if both ASEs adopted an ASEL fate (Fig. S2C-D). Symmetrizing the cell fate of postsynaptic AWC neurons had no effect on ASE>AWC connectivity (Fig. S2C-D). Collectively, these findings show that presynaptic cell fate is important, but not sufficient to establish left biased ASE>AWC connectivity, whereas postsynaptic cell fate does not appear to be important for connectivity (Fig. S2C-D). These observations further imply that the olfactory functions of AWC neurons may not be affected by the inclusion of AWC into the ASE salt-sensing circuit. Consistent with this interpretation, olfaction of AWC-sensed odors remained unchanged over a wide range of concentrations of different odorants between naïve and conditioned animals (Fig. S3A). Lastly, these genetic observations imply factors other than cell fate determinants that influence synaptic connectivity, as asymmetry persists even when ASE cell fate is symmetrized.

### The insulin-signaling pathway regulates ASE>AWC synaptic connectivity

Since asymmetric fate of ASE was insufficient to establish asymmetric connectivity between ASE and AWC neurons, we searched for potential signaling pathways that may be involved. Since the insulin signaling pathway is crucial for *C. elegans* salt sensation and learning [23-28], we tested whether the ASE>AWC synaptic connection is regulated by insulin-signaling. We found that a mutation in the sole insulin/insulin-like growth factor receptor (IIR) in *C. elegans, daf-2* [29], or in *age-1*/PI3K, which encodes a phosphatidylinositol-4,5-bisphosphate 3-kinase subunit that functions downstream of the daf-2/IIR [30] (Fig. 3B) resulted in a symmetrized ASE>AWC synaptic connection (Fig. 3A, C-D, Fig. S4A-B). The FoxO like transcription factor *daf-16*/FoxO is inhibited by *daf-2*/IIR signaling via phosphorylation and subsequent translocation of the transcription factor from the nucleus to the cytosol [31](Fig. 3B). We therefore pan-neuronally expressed an unphosphorylatable, constitutively active DAF-16(S4A) mutant cDNA, which is not inhibited by daf-2/IGR [32]. This manipulation should mimic loss of *daf-2*/IGR function, and consistent with this expectation, we found ASE>AWC synaptic connectivity to be symmetrized (Fig.3C-D). Collectively, these findings show that insulin signaling is required to establish asymmetric synaptic connectivity.

**Fig. 3.**
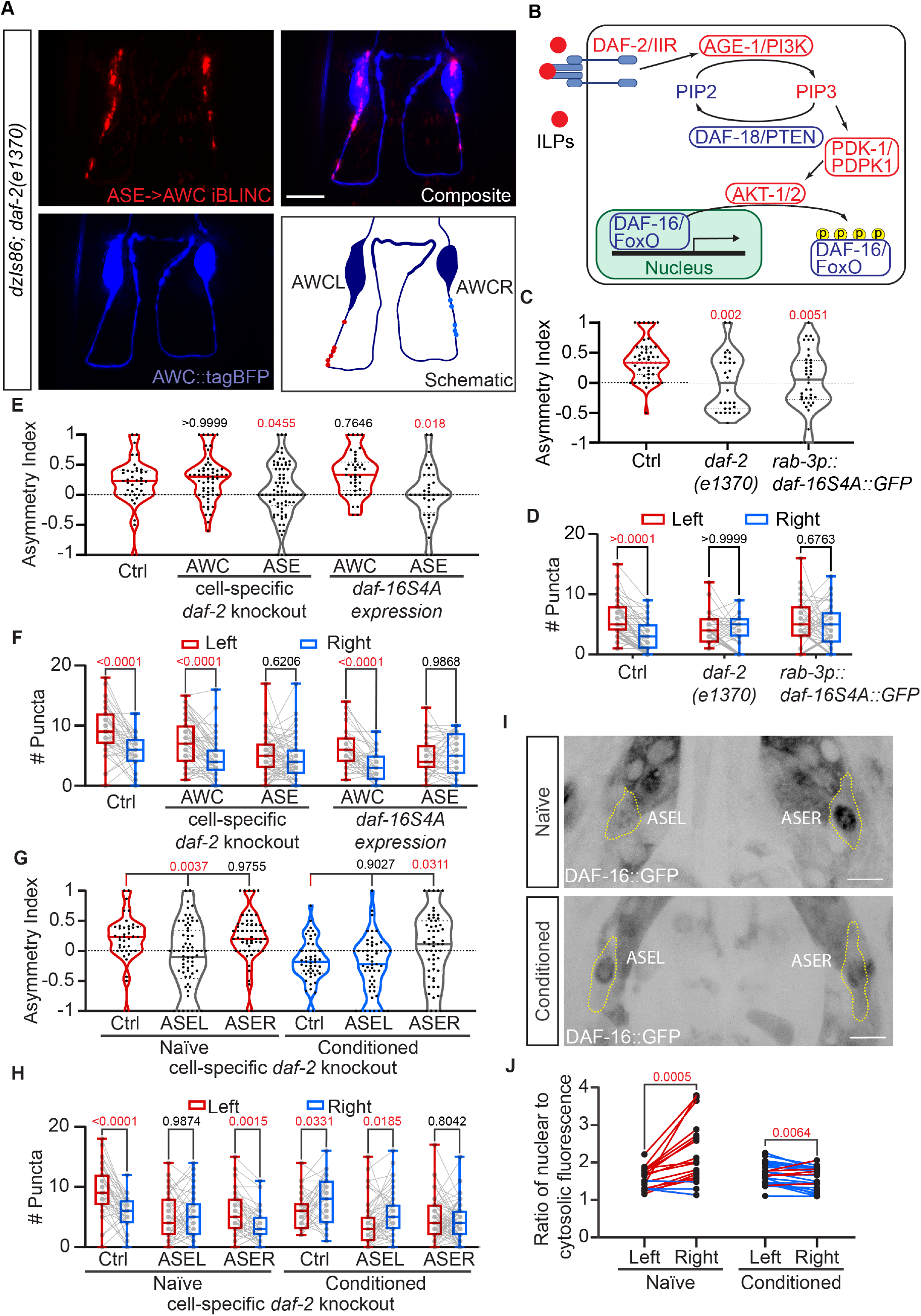
Asymmetric insulin signaling in ASE presynaptic neurons establishes lateralized ASE>AWC connectivity. **(A)** Representative images of ASE>AWC iBLINC signal in red and AWC cell bodies in blue in a *daf-2(e1370)* mutant animal. Scale bar = 10 μm. In the schematic, red and blue puncta indicate location of iBLINC puncta on the left and on the right respectively. **(B)** Diagram showing the insulin signaling pathway with *C. elegans* and vertebrate gene names indicated. (**C-D**). Asymmetry indices (C) and number of puncta (D) comparing iBLINC in wildtype (N = 53), *daf-2(e1370)* mutants (N = 31), and pan-neuronal expression of overactive DAF-16S4A (n=39). (C) was analyzed with unmatched one-way ANOVA. P value from Tukey multiple comparison test against control are labeled on graph. (D) was analyzed with two-way matched ANOVA with Sidak’s multiple comparison. (**E-F**) Asymmetry indices (E) and number of puncta (F) showing iBLINC of the *daf-2* floxed allele (Ctrl) and cell specific knockouts through expression of Cre recombinase (N = 40 for control, 57 for AWC, and 74 for ASE) or expression of constitutively active DAF-16S4A (N = 40 for AWC, 36 for ASE. (E) was analyzed with unmatched one-way ANOVA, P value from Tukey multiple comparison test against control are labeled on graph. (F) was analyzed with two-way matched ANOVA with Sidak’s multiple comparison. (**G-H**) Asymmetry indices (G) and number of puncta (H) showing iBLINC of the *daf-2* floxed allele (Ctrl) and cell specific knockouts of *daf-2* in naïve animals (N = 40 for control, 57 for ASEL, and 66 for ASER) or conditioned animals (N = 49 for Control, 57 for ASEL and 57 for ASER). (G) was analyzed with unmatched one-way ANOVA. P value from Tukey multiple comparison test against control within conditioning group are labeled on graph. (H) was analyzed with two-way matched ANOVA with Sidak’s multiple comparison. (**I-J**) Representative image of DAF-16::GFP endogenous reporter in ASE (I) and quantification of nuclear to cytosolic ratio of average DAF-16::GFP signal (J) in naïve (N = 18) and conditioned (N = 22) animals, plotted as a line graph with each line as one animal. Scale bar = 10 μm. (J) was analyzed with two-way matched ANOVA with Sidak’s multiple comparison. Red and blue lines indicate animals with a higher nuclear-to cytosol ratio in ASER and ASEL respectively.

To interrogate whether insulin signaling is functioning in presynaptic ASE or postsynaptic AWC neurons to direct synaptic connectivity, we created a floxed allele of *daf-2*/IIR by inserting loxP sites flanking the third and fourth exon, which upon recombination should remove all *daf-2*/IIR isoforms with extracellular domains (Fig. S5A). Cell-specific expression of Cre recombinase in presynaptic ASE but not in postsynaptic AWC neurons symmetrized connectivity like in genomic *daf-2*/IIR loss of function mutant animals (Fig. 3E-F), indicating that *daf-2*/IIR functions in ASE to regulate ASE>AWC connectivity. Similar results were obtained when we expressed the unphosphorylatable DAF-16(S4A) mutant in ASE and AWC, respectively (Fig. 3E-F). The change in ASE>AWC connectivity appeared independent of the DAF-2c/IIR splice variant and the CASY-1/calsyntenin cell adhesion molecule (Fig. S4A-B), both of which are important for signaling at ASER synapses during salt aversive learning [24]. We conclude that insulin signaling in ASE directs ASE>AWC connectivity through a CASY-1-independent pathway.

Interestingly, genetically removing *daf-18*/PTEN, a negative regulator of the insulin signaling pathway [33] (Fig. 3B), also resulted in symmetrization of ASE>AWC connectivity (Fig. S4A-B). Thus, both global inactivation and over-activation of the insulin signaling pathway results in symmetrized ASE>AWC connectivity, raising the possibility that asymmetric insulin signaling in ASEL and ASER neurons regulates synaptic connectivity. To test this hypothesis, we knocked out *daf-2*/IIR through cell-specific expression of Cre in ASER and ASEL, respectively. We observed that in naïve animals, removal of *daf-2*/IIR in ASEL induces symmetrization through a reduction of connectivity on the left. Conditioned animals in which *daf-2*/IIR was knocked down in ASEL, however, still changed to right-side biased connectivity. In contrast, knockout of *daf-2*/IIR in ASER in naïve animals had no effect on connectivity but resulted in symmetrization of ASE>AWC connectivity in conditioned animals (Fig. 3G-H). These findings suggest that DAF-2/IIR serves the same function in ASEL and ASER, i.e. induce the formation of new synapses, and that the DAF-2/IIR insulin receptor is asymmetrically activated. To directly test this hypothesis, we measured the level of insulin pathway activation in ASE neurons through an endogenously tagged DAF-16::GFP translational reporter [34] (Fig. 3I). The DAF-16/FoxO transcription factor translocates from the nucleus to the cytosol in response to insulin pathway activation [31]. We found that in naïve animals, the nuclear-to-cytosolic ratio of DAF-16::GFP fluorescence is lower in ASEL than in ASER neurons, while the reverse is true in conditioned animals (Fig. 3J). Therefore, the insulin pathway is more active in ASEL in naïve animals, and more active in ASER in conditioned animals, correlating with ASE>AWC asymmetric synaptic connectivity. Together, these findings suggest that asymmetric insulin signaling is superimposed on the asymmetric cell fates of ASE neurons to establish plastic, asymmetric synaptic connectivity between ASE and AWC neurons. We propose that activation of the insulin pathway induces formation of synapses between ASE and AWC neurons while inactivation may reduce connectivity.

### The insulin-like peptide INS-6/ILP regulates ASE>AWC non-cell-autonomously in a paracrine manner

The *C. elegans* genome encodes at least 40 insulin-like peptides (ILP) [35]. Previous work had suggested a role for the insulin-like peptide *ins-6*/ILP in response to large increases in salt stimuli, where it had been proposed that INS-6/ILP functions from ASE to modulate the AWC response to changes in salt concentration [23]. We therefore tested whether loss of *ins-6*/ILP affected ASE>AWC synaptic connectivity. We found that *ins-6*/ILP null mutants displayed a right-side bias in ASE>AWC synaptic connectivity, unlike *daf-2*/IIR mutant animals (Fig. 4A-C, cf. Fig. 3B-C). Since DAF-2/IIR is the presumed sole ILP receptor, this shows that while *ins-6*/ILP is an important regulator of ASE>AWC connectivity, other ILPs are likely involved. Consistent with this interpretation *ins-6*; *daf-2* double mutant animals exhibited the same symmetrized phenotype as *daf-2*/IIR mutant animals alone (Fig. S4C-D). This notion is further supported by the observation that mutants that disrupt processing or secretion of all ILPs, such as *egl-3*/PC2 [36]) and *unc-31*/CAPS [37], respectively, also show symmetrized synaptic connectivity between ASE and AWC neurons (Fig. S4C-D).

**Fig. 4.**
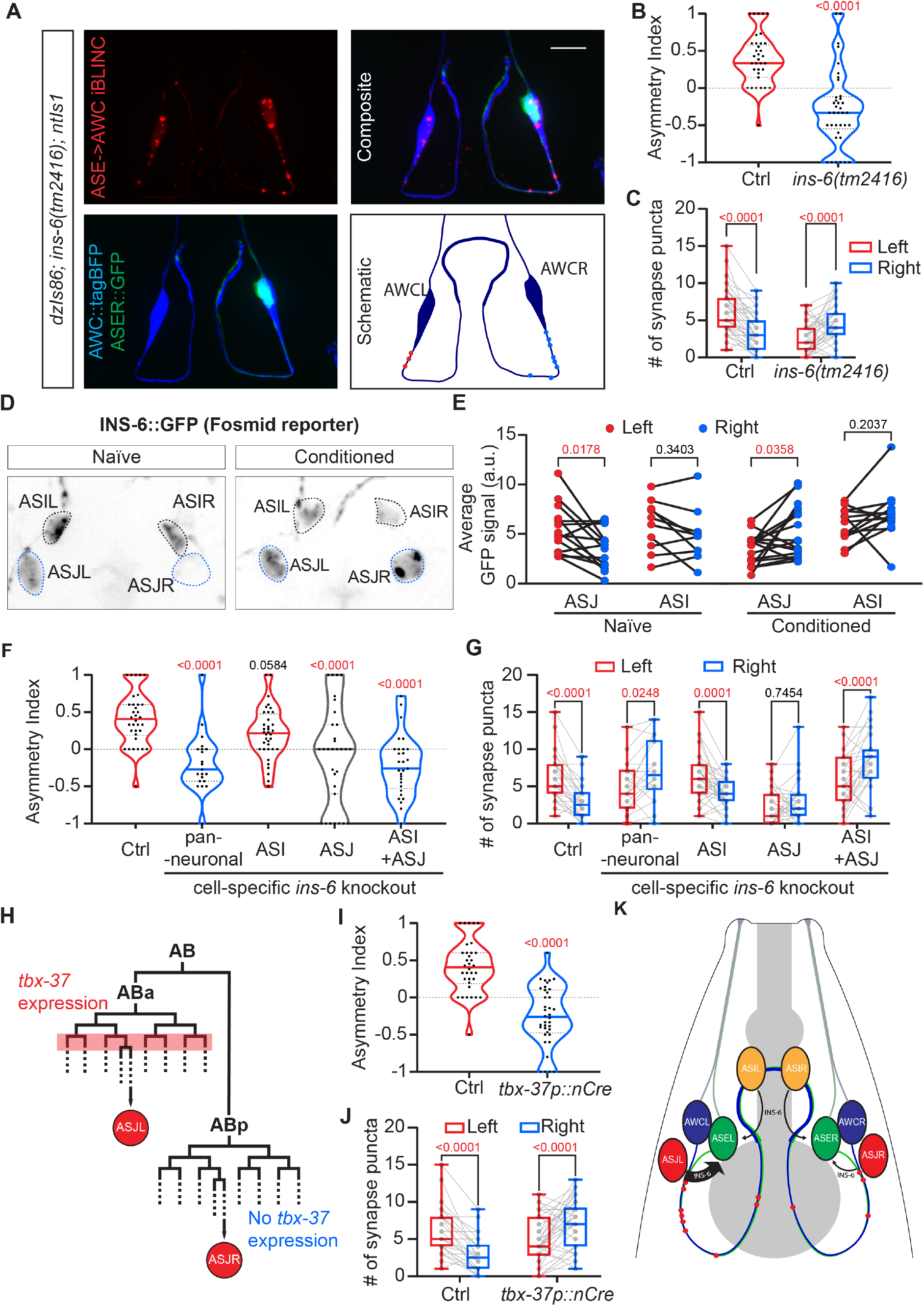
Asymmetric, paracrine *ins-6*/ILP signaling establishes lateralized ASE>AWC connectivity. (**A**) Representative images of ASE>AWC iBLINC in red and AWC in blue in *ins-6(tm2416)* mutant animals. Scale bar = 10 μm. In the schematic, red and blue puncta indicate location of iBLINC puncta on the left and right respectively. (**B-C**) Asymmetry indices (B) and number of puncta (C) comparing iBLINC in wildtype (N = 53) and *ins-6(tm2416)* mutants (N = 37). (B) analyzed with two-tailed unpaired t test. (C) was analyzed with two-way matched ANOVA with Sidak’s multiple comparison. **(C)** Representative images of an INS-6::GFP translational reporter in naïve and conditioned animals, showing expression of INS-6 in ASI and ASJ. Scale bar = 10 μm **(E)** Quantification of GFP signal in ASI and ASJ in naïve and conditioned animals, plotted as a line graph with each line as one animal. Analyzed with two-way matched ANOVA with Sidak’s multiple comparison. (**F-G**) Asymmetry indices (F) and number of puncta (G) showing iBLINC of the *ins-6*/ILP floxed allele (Ctrl) and cell specific knockouts through expression of Cre recombinase (N = 34 for control, 21 for pan-neuronal KO, 36 for ASI KO, 31 for ASJ, and 29 for ASI+ASJ KO). (F) analyzed with unmatched one-way ANOVA. P value from Tukey multiple comparison test against control are labeled on graph. (G) analyzed with two-way matched ANOVA with Sidak’s multiple comparison. (**H**) Schematic illustrating the cell lineage giving rise to ASJL or ASJR, with transient expression of tbx-37 in the ABa but not ABp lineage indicated. (**I-J**) Asymmetry indices (I) and number of puncta (J) showing iBLINC of the *ins-6*/ILP floxed allele (Ctrl) and ASJL-specific knockout through expression of Cre recombinase under the tbx-37 promoter (N = 34 for control, (N = 38) for *tbx-37p*). (I) analyzed with two-tailed unpaired t test. (J) analyzed with two-way matched ANOVA with Sidak’s multiple comparison. (**K**) Model of INS-6 regulating ASE>AWC connectivity through paracrine action in wild type naïve animals.

To determine in which tissue *ins-6*/ILP functions and how it is regulated, we conducted transgenic rescue experiments. Expression of *ins-6*/ILP under its own promoter, but not under heterologous promoters driving expression in ASE either bilaterally or individually could fully rescue the changes in ASE>AWC connectivity in *ins-6*/ILP mutants (Fig. S5E-F), suggesting that *ins-6*/ILP expression may not function in ASE neurons or is specifically regulated. We therefore analyzed animals transgenically expressing an INS-6::GFP protein fusion from a fosmid-based reporter, which should contain most if not all regulatory elements controlling expression of *ins-6*/ILP. Consistent with previous studies of transcriptional *ins-6p* promoter fusions with GFP as well as single cell transcriptomic studies [36, 38, 39], we found expression primarily in the two pairs of ASI and ASJ sensory neurons (Fig. 4D). Intriguingly, the INS-6::GFP reporter exhibited asymmetric expression in ASJ neurons of naïve animals with a higher GFP signal in the left ASJL(left) neuron compared to the ASJR(right) neuron, but symmetric expression in both ASI neurons (Fig. 4D-E). Moreover, conditioned animals changed asymmetric expression of the INS-6::GFP reporter in ASJ from a left-sided bias to a right-sided bias, without obviously affecting symmetric expression of the reporter in ASI neurons (Fig. 4D-E).

The positive correlation of asymmetric expression of the INS-6::GFP insulin-like peptide in ASJ neurons with the asymmetry of the ASE>AWC synaptic connection in naïve animals after high salt conditioning, suggested that control of ASE>AWC synaptic connectivity required *ins-6*/ILP expression from ASJ. To test this hypothesis, we introduced flanking *LoxP* sites in the *ins-6*/ILP locus (Fig. S5A) and expressed Cre recombinase to remove *ins-6*/ILP in a cell-specific manner. Pan-neuronal expression of Cre recombinase recapitulated the right-side biased asymmetry in the ASE>AWC connectivity of the *ins-6*/ILP null mutant (Fig. 4F-G, cf. Fig. 4B). In contrast, removing *ins-6*/ILP from ASI neurons had no effect on ASE>AWC asymmetric connectivity (Fig. 4F-G), likely because the asymmetric source of INS-6/ILP in ASJ neurons was still present. Removing *ins-6* from ASJ symmetrized the ASE>AWC connectivity, because now only the symmetric source of INS-6/ILP in ASI neurons remained. Only when *ins-6*/ILP was removed in both ASI and ASJ neurons, we observed the change to right-side bias in the ASE>AWC connectivity that is characteristic of complete loss of *ins-6* function (Fig. 4F-G). Taken together, these findings showed that while *ins-6*/ILP asymmetric expression from ASJ is an important factor for asymmetric ASE>AWC connectivity, *ins-6*/ILP expression in ASI neurons is important as well. We propose that both expression levels and spatial origin of *ins-6*/ILP are crucial for regulating ASE>AWC connectivity, where *ins-6*/ILP acts locally as a paracrine signal.

One prediction of this model is that removal of *ins-6*/ILP unilaterally on the left would result in ASE>AWC connectivity to exhibit a right-side bias. There is no known promoter that expresses exclusively only in ASJL or ASJR to allow for asymmetric expression of Cre recombinase. However, the ASJL/R neurons originate from the ABa and ABp cell lineage branches, respectively, which are distinguished by transient expression of the TBX-37/T-box transcription factor during early embryonic stages in the ABa lineage but not in the ABp lineage [40] (Fig. 4H). When we expressed Cre recombinase under control of the *tbx-37p* promoter to selectively remove *ins-6*/ILP in the ABa lineage, which gives rise to ASJL, we observed a right-sided bias in ASE>AWC connectivity (Fig. 4I-J). These findings suggest that asymmetric expression of *ins-6*/ILP on the left is necessary to establish left-biased asymmetric ASE>AWC connectivity in naïve animals.

### Restructuring the salt sensing circuit results in predictable changes in behavior

Our studies established a correlation between synaptic connectivity and an associative salt learning paradigm. Left-sided connectivity of the ASE>AWC module favored preference of the naïve conditions, whereas right-sided connectivity due to prior conditioning at elevated salt concentrations resulted in a preference for higher salt concentrations. Since the change in connectivity between ASE and AWC sensory neurons in response to salt is mediated by ILP signaling, and loss of the INS-6/ILP results in a right-sided bias of the ASE>AWC module, we asked whether genetic removal of *ins-6*/ILP resulted in changes in behavior. We found that naive *ins-6*/ILP mutant animals preferred 100 mM NaCl rather than 33 mM NaCl, similar to wild type animals that had been conditioned on 100 mM NaCl (Fig. 5A). Conditioning *ins-6*/ILP mutant animals with NaCl for 12 hours had no measurable additional effect on salt preference or connectivity (Fig. 5A, Fig. S4G-H). Therefore, a genetic manipulation of the ASE>AWC module that led to the same change in network architecture as conditioning on high salt for 12 h had identical effects on salt preference in the associative learning paradigm. These findings are consistent with the conclusion that a right-side bias in ASE>AWC network architecture is sufficient for salt preference.

**Fig. 5.**
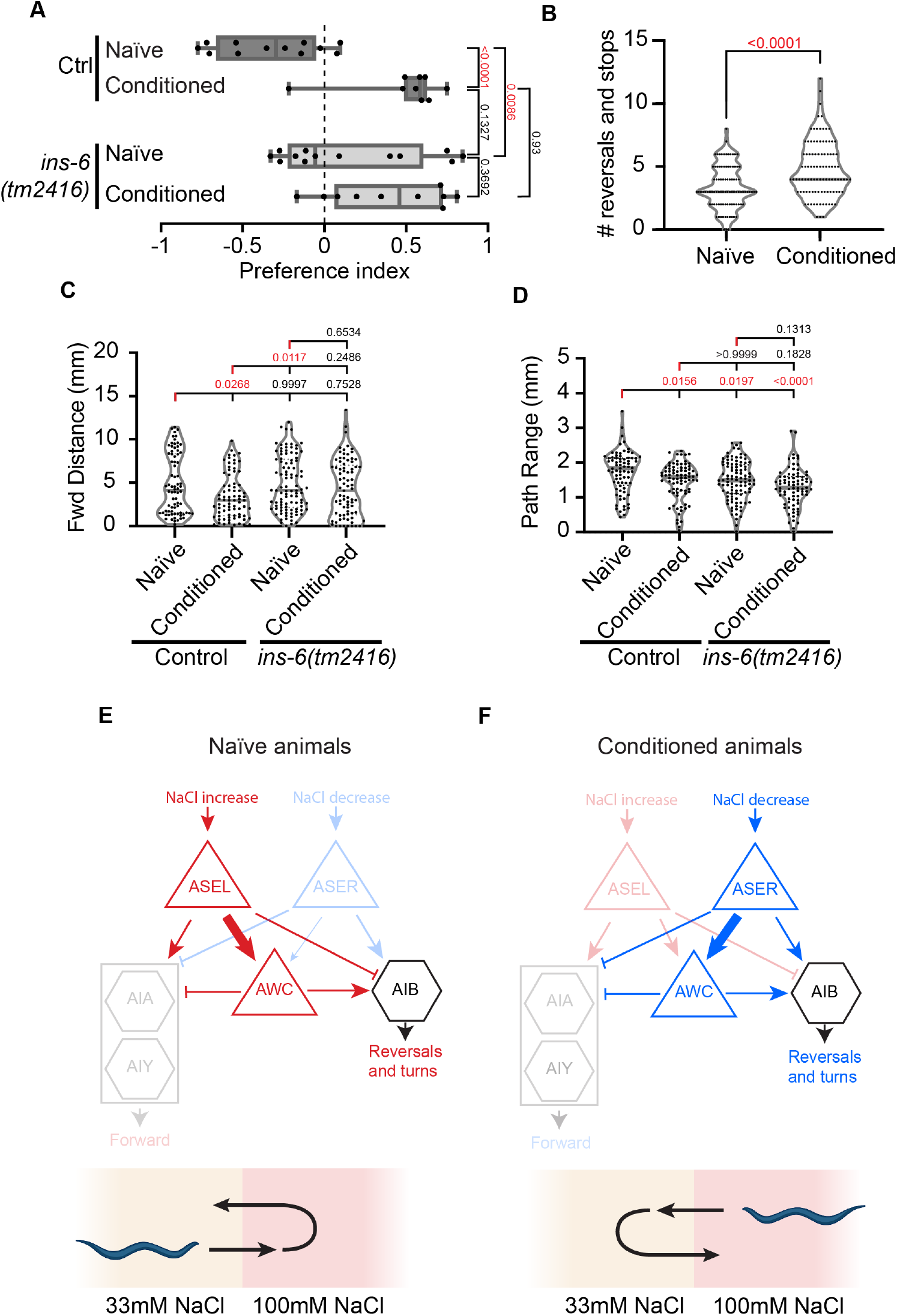
Changed asymmetric ASE>AWC connectivity predicts behavioral changes. **(A)** Preference indices of wildtype or *ins-6(tm2416)* mutant animals subjected to the same conditioning protocol and choice assay as described in Fig.1A. N = 12 and 9 for control naïve and conditioned trials respectively, N = 13 and 10 for *ins-6* mutant naïve and conditioned animals respectively. Analyzed with one-way ANOVA **(B)** Number of stops and reversals scored for 30 seconds on 33mM NaCl plates with naïve (N = 95) or conditioned (N = 99) animals. Analyzed with unpaired t-test. (**C-D**) Wildtype or *ins-6(tm2416)* mutant animals are subjected to the conditioning protocol, then subjected for 30 seconds to high resolution worm tracking using the Multi-Worm tracker v1.5.3a. Forward distance (C) and path range (D) are plotted here. Forward distance is defined by the total distance animals have traveled forward regardless of orientation. Path range is defined by the distance between the final location of the animal, and the geographic center of the path traveled. Analyzed with one-way unmatched ANOVA with Tukey multiple comparison. (**E-F**) Simplified circuit diagram of the salt-sensing module in *C. elegans*. In naïve animals (E), the left ASE>AWC connection is stronger than the right ASE>AWC connection, whereas the opposite is true in conditioned animals (F). Therefore, AWC is more likely activated by salt increases in naïve animals and salt decreases in conditioned animals, respectively. This results in an increased probability of turns and reversals in naïve versus conditioned animals upon encountering, respectively, acute increases versus decreases in salt concentrations.

We next asked whether the observed differences in performance of the learning paradigm are reflected in changes in locomotory behavior. *C. elegans* uses two general locomotory strategies for chemotaxis: klinokinesis and klinotaxis [41, 42]. While klinotaxis relies on small, continuous changes to orientation, klinokinesis is achieved by more drastic changes in orientation (pirouettes, reversals) that lead to changes in direction. We therefore first measured the number of stops and reversals in naïve animals and animals conditioned on 100 mM NaCl for 12 h when placed on plates with the naïve 33 mM NaCl concentration. We found a significant increase in stops and reversals in conditioned versus naïve animals when placed on the naïve salt concentration (Fig. 5B), suggesting that the restructured sensory circuit changes its klinokinetic strategy in search of preferential salt concentrations. We next used a single-worm tracking approach to establish high resolution, quantitative locomotory parameters in response to acute changes in environmental salt concentration [43]. We found several behavioral parameters unaffected by either salt conditioning or in *ins-6*/ILP mutants (Fig. S6), including path curvature, which would be expected to change if klinotaxis was primarily affected, as well as forward time, pause time, and dwelling time, among others. In contrast, path range as a measure of area explored by a worm in a given time frame, was reduced in both salt-conditioned and *ins-6*/ILP mutant animals (Fig. 5C). Therefore, salt-conditioned and *ins-6*/ILP mutant animals explore less when placed on the naïve (33 mM) NaCl concentration. The reduced exploration was not the result of less forward movement, because *ins-6*/ILP mutant animals did not show reduced forward motion, but rather due to increased reversals and turns (Fig. 5D). However, salt conditioned animals did display reduced forward motion and conditioned *ins-6*/ILP mutant animals showed some defects in backward movement (Fig. 5D, Fig. 6E-F), suggesting possibly additional differences in circuit architecture between salt conditioned and *ins-6*/ILP mutant animals.

ASEL, ASER and AWC sensory neurons have been shown to form synapses onto the interneurons AIA, AIB and AIY [16]. ASEL forms excitatory synapses onto AIA and AIY and inhibitory synapses onto AIB [44], while both ASER and AWC form excitatory synapses onto AIB and inhibitory synapses onto AIA and AIY [45, 46]. AIA and AIY induce forward motion while AIB promotes reversals and turns [44]. Our results together with these studies suggest a salt sensing and learning circuit (Fig. 5E-F), where in naïve animals (E), the ASEL>AWC synaptic connectivity is stronger than ASER>AWC connectivity, whereas in conditioned animals the converse is true (F). We propose that in naïve animals, AWC is more likely to respond to salt increases as opposed to salt decreases. A response in AWC may lead to activation of AIB and inhibition of AIA and AIY, thereby dampening the activation of AIA and AIY and inhibition of AIB generated by the direct synapses from ASEL. This would result in an increased chance of reversal and decreased chance of forward motion when encountering a higher salt environment with a net effect of a preference for lower salt. In contrast, the converse is true in conditioned animals. AWC now amplifies signals from ASER, which leads to an increased chance of reversal and a reduced chance of forward motion in response to decreases in salt concentration, with a net effect of a preference for higher salt concentrations. It should be noted that changes in connectivity are unlikely to be the sole mechanisms for salt learning. For example, the modulation of glutamatergic transmission between the ASER and AIB interneurons is an important component of salt learning, with mechanisms that include both the ASER presynaptic side and the postsynaptic AIB side [47, 48].

In summary, we have shown that the *C. elegans* connectome is plastic, i.e. that network architecture is not fixed but can be changed by experience. The resulting changes correlate with predictable differences in behavior. The interindividual variability we observe is likely not genetically encoded, because the progeny of an animal displays the same range of variability in connectivity regardless of the connectivity of the parent (Fig. S2E). We propose that this variation does also not result only from stochastic circuit development, as documented in Drosophila [49]. Instead, the plasticity of the salt sensing circuit is mediated by asymmetric insulin-like signaling in an experience-dependent manner. The widespread yet specific expression of insulin-like peptides in the nervous system not only in worms [27], but also across phyla [50], raise the possibility of conserved functions in mediating changes in network architecture of mammalian connectomes. In this context it is interesting to note that both insulin-like peptide signaling, and asymmetric brain functionality have been implicated in neuropsychiatric conditions such as autism spectrum disorders and schizophrenia [51, 52].

## Acknowledgments

We thank S. Emmons, O. Hobert, A. Jenny, and P. Kurshan as well as members of the Bülow laboratory for comments on the manuscript and discussions throughout the course of this work. We are grateful to O. Hobert, D. Kim and P. Kurshan for reagents, and the Caenorhabditis Genetics Center for some of the strains used in this study. We acknowledge help with imaging from the Analytical Imaging Facility at Albert Einstein College of Medicine

## Author contributions

Conceptualization: LTHT, GAL, SJC, HEB

Methodology: LTHT, GAL, SJC, JH, HEB

Investigation: LTHT, GAL, SJC, JH, CCP

Visualization: LTHT, GAL, HEB

Funding acquisition: LTHT, HEB

Project administration: HEB

Supervision: LTHT, GAL, HEB

Writing – original draft: LTHT, GAL, HEB

Writing – review & editing: LTHT, GAL, HEB

## Data and materials availability

All data are available in the main text or the supplementary materials

## Supplementary Data

### Materials and Methods

#### C. elegans strains and genetics

All C. elegans strains were grown on King’s agar plates with E. coli (OP50) as a food source as described [53], at 20°C unless otherwise specified. Strains used in this work can be found in Table S1.

#### Molecular Biology

Cloning of all constructs were carried out using standard molecular biology methods. A list of primers used can be found in Table S2. For a list of plasmids used and detailed generation methods of constructs see Table S3 and S4.

#### Transgenesis

All extrachromosomal arrays were generated by injecting a mixture of the required plasmids along with pBluescript to a final concentration of 100 ng/μl of DNA. A detailed list of all the extrachromosomal transgenic lines generated can be found in Table S5. Integration of arrays were performed with a standard gamma-ray radiation protocol. All integrated arrays were backcrossed four times prior to further use.

For CRISPR/Cas9 mediated genome editing, after appropriate guide RNA sites were selected using the IDT Cas9 crRNA design tool, oligos in the form of TAATACGACTCACTATA(gRNA)GTTTTAGAGCTAGAAATAGCAAG were ordered, where (gRNA) is the 20nt of the guide RNA sequence before the PAM motif, optimized for T7 promoter transcription. These oligos were used in PCR reactions as a forward primer in conjunction with the reverse primer AAAAGCACCGACTCGGTG (oLT337) to generate a sgRNA transcription template from pDD162 (containing the sequence of the tracrRNA). sgRNA was then transcribed from this template using HiScribe T7 in-vitro transcription kit (NEB) following the manufacturer’s instructions and purified with Monarch RNA cleanup kit (NEB).

The resulting sgRNA was used in an injection mix at 20 ng/μl per each sgRNA, together with 250 ng/μl of Alt-R Cas9 endonuclease (IDT) and repair templates (100 ng/μl for single-strand oligos, 200 ng/μl for PCR products or plasmids). Injection and CRISPR efficiency were monitored through Co-CRISPR strategy with *dpy-10(cn64)* or *unc-58(e665)* conversions, or through *rol-6(su1006)* co-injection markers. A list of all genome-edited strains and detailed methodology can be found in Table S6.

#### NaCl conditioning protocol

Populations of animals were synchronized by egg prep with a NaOH and bleach solution and then grown on King’s Agar plats at 20° C for 3 days. Animals were continuously well fed for the duration and for at least two generations prior to egg prep. The animals were then washed in 1 mL of M9 buffer and collected in a microcentrifuge tube. Worms were allowed to settle before the supernatant was removed and were washed again with water. Fifty to one-hundred animals were then placed on King’s Agar plates as control or King’s agar with modifications as indicated. All plates were seeded with 300 μl of OP50 grown O/N in 2xYT media at 37° C prior. Animals were kept at 20° C for the time indicated for each experiment.

#### Salt preference assay

Assay plates were generated on quadrant petri dishes (VWR, #25384-348). 16ml of King’s agar liquid with either 33mM NaCl or 100mM NaCl were deposited into each quadrant to create the assay plates as shown in Fig.1a. After the agar solidified and dried, the plates were used within a week. Animals were conditioned either on King’s Agar (with 33mM NaCl) or high salt King’s Agar plates (with 100mM NaCl) as described above. Immediately before the assay, a 2% agar solution was pipetted between the quadrants but not at the center to create bridges between the quadrants. The animals were collected with M9 buffer and washed once in water before placed at the center of an assay plate. The animals were then allowed to move for 15 min at room temperature before animal distribution was recorded by digital photography. The number of animals on each quadrant was manually counted in imageJ and the preference index was calculated using the following formula:

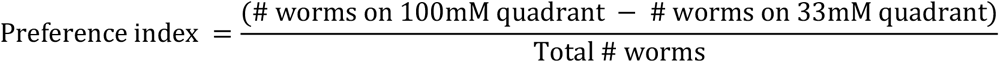

#### Behavioral tracking

Animals were conditioned at 100 mM NaCl for 12 hours as described above. Animals were then single-picked from control or conditioning plates and placed on an assay plate containing King’s Agar without food. After a thirty second acclimatization period, behavior was recorded using a DinoLite AM413T camera with a 630 nm red-light source in a setup as described in[54]. Videos were recorded at a resolution of 1280 × 720 pixels with a field of view of 2.5 cm for thirty seconds. Immediately after recording and, therefore, blind to behavioral analysis, animals were mounted in 1 mM levamisole on 5% agarose pads and ASEL/R puncta number was quantified using the eyepiece of a Plan-Apochromat 63x/1.4 objective on a Zeiss Axioimager.

All videos were analyzed using TierPsy Multi-Worm tracker v1.5.3a[43], which is available for download from https://github.com/Tierpsy/tierpsy-tracker. TierPsy Features were extracted for each video and analyzed in Prism 9 using an unpaired t-test between groups for each feature. The details of each behavioral feature are described here: https://github.com/ver228/tierpsy-tracker/blob/master/docs/OUTPUTS.md.

#### Quantification of connection asymmetry

Strains containing either *dzIs86* (ASE>AWC iBLINC)[16] or *dzIs127* (*flp-6p::cla-1::gfp*, this study) were used to visualize ASE to AWC synapses and ASE pre-synaptic active zones, respectively. Young adults were immobilized using 1 mM levamisole and mounted on 5% agarose pads. The number of puncta on each side was then counted manually on the eyepiece using a Plan-Apochromat 63x/1.4 objective on a Zeiss Axioimager. The asymmetry index was calculated for each individual animal using the following formula:

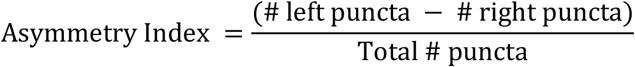

#### Fluorescent microscopy and quantification

INS-6::GFP quantification: Animals expressing (*ins-6[30042]::S0001_pR6K_Amp_ 2xTY1ce _EGFP_ FRT_rpsl_neo_FRT_3xFlag*)[55] were conditioned for 12 hours on 100 mM NaCl plates and mounted for imaging using a Plan-Apochromat 63x/1.4 objective on a Zeiss Axioimager. Transgenic control and conditioned animals were imaged with the only selection criteria being a dorsal-ventral mounted orientation. All animals were imaged in a z-stack to capture the cell bodies of all GFP+ neurons using the same parameters across experiments. Images were analyzed in imageJ to calculate the fluorescence by defining the cell bodies of ASJL/R and ASIL/R as a ROI. Corrected total cell fluorescence was calculated as the (integrated density of the ROI - (area of ROI x mean background fluorescence)).

ASE DAF-16::GFP nuclear localization: Animals containing *daf-16(ot971[daf-16::GFP]) I; dzEx2181 [flp-6p::tagBFP + flp-6p::nls::mCherry])* were immobilized using 1 mM levamisole and mounted on a 5% agarose pad, then imaged under a Plan-Fluor Nikon 100x/1.4 objective on a Nikon CSU-W1 Spinning Disk Confocal with identical imaging parameters and appropriate z-stack. Only animals that were dorsal-ventrally oriented were imaged. Using the threshold and particle analysis tool in ImageJ, the tagBFP and mCherry channel were used to define the ASE cell body and nucleus ROI respectively for each relevant Z slides. The average GFP signal in the nucleus was then calculated as the (total GFP signal sum in the nuclear ROI) / (area sum of nuclear ROI) area across relevant Z slides. The average GFP in the cytosol was calculated by (total GFP signal sum in cell body – total GFP signal sum in nucleus) / (area sum of cell body ROI – area sum of nuclear ROI) across relevant Z slides. The nuclear localization ratio was then obtained as (average nuclear GFP signal) / (average cytosolic GFP signal).

#### Statistical Analysis

All statistical analyses were performed using the Prism 9 Statistical Software suite (GraphPad). Left-right asymmetric comparisons are performed using matched two-way ANOVA with post hoc Sidak’s multiple comparison test. All comparisons of means are analyzed with Student t-test or one-way ANOVA with Tukey correction where applicable. Throughout the manuscript, asymmetry indices are color coded in red for significantly left-biased, blue for significantly right biased, and grey for no significant bias. All box and whisker graphs indicate ranges, quartiles, and medians, overlayed with individual data points graphed as a line for each animal. P values are indicated in red (if <0.05) or black (if >0.05).

### Supplementary Figures

**Fig. S1.**
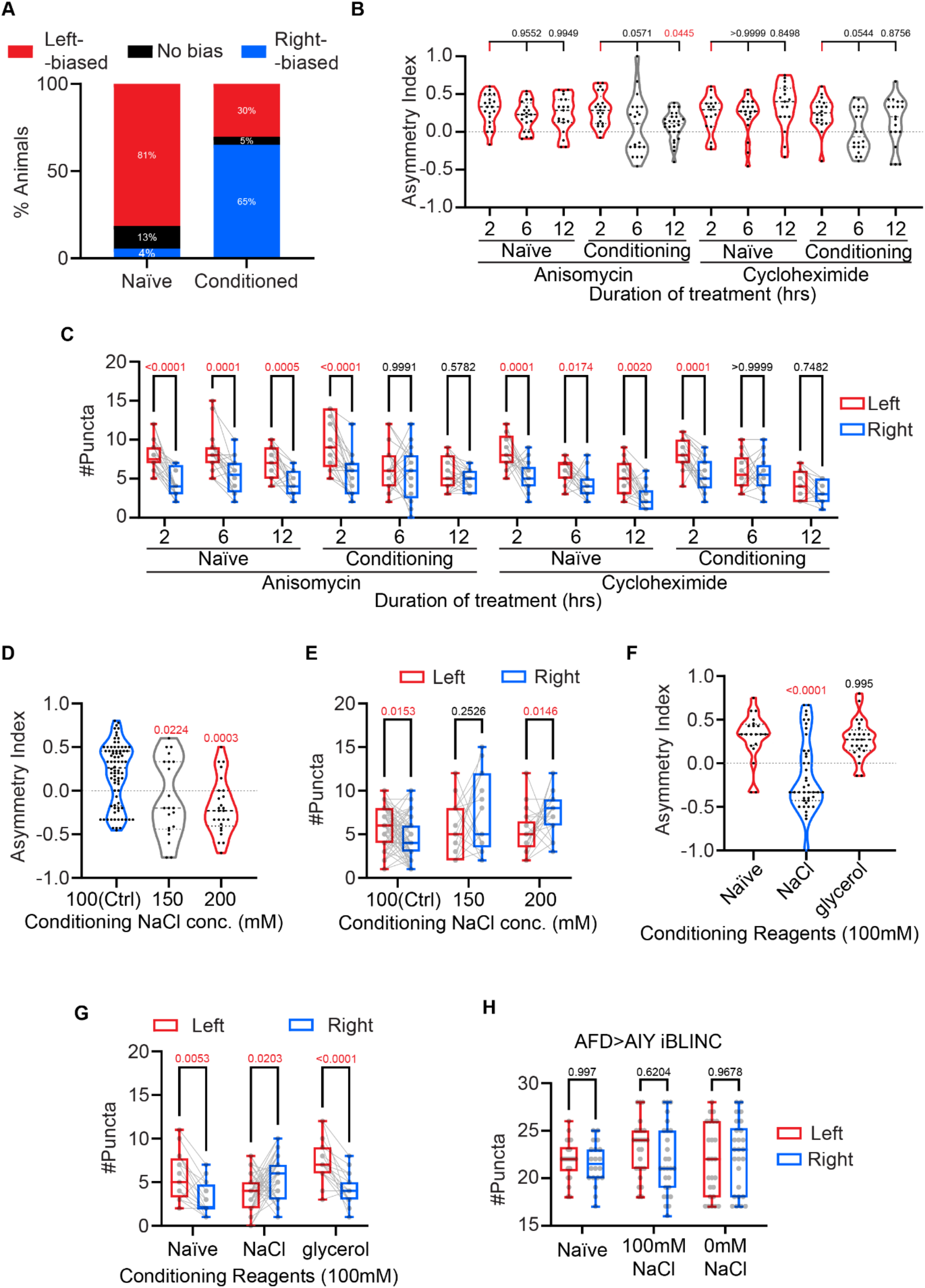
(**A**) The percentage of animals grouped as left-biased (More synapses on the left than right), no bias (Synapse number is the same in left and right), and right-biased (Less synapses on the left than right), plotted as a bar graph for both naïve (N = 54) and conditioned (N = 43) animals. Percentage of the population of each category are labeled on the bar. (**B-C**) Asymmetry indices (B) and number of puncta (C) for ASE>AWC iBLINC under anisomysin treatment with or without simultaneous 100mM conditioning (N = 20, 20 and 19 for 2, 6 and 12 hours without conditioning respectively, and N = 21, 21 and 27 for 2, 6 and 12 hours with conditioning respectively), and under cycloheximide treatment with or without simultaneous 100mM conditioning (N = 17, 19 and 17 for 2, 6 and 12 hours without conditioning respectively, and n =18, 20 and 19 for 2, 6 and 12 hours with conditioning respectively). (B) analyzed with two-tailed unmatched t-test. (C) was analyzed with two-way matched ANOVA, with p value from Sidak’s multiple comparison test labeled on graph. (**D-E**) Asymmetry index (D) and number of puncta graphs (E) showing iBLINC after animals are raised in 100mM NaCl and conditioned for 12 hours with the indicated concentration of NaCl. N = 80 for 100mM, 21 for 150 mM, and 25 for 200mM. (D) was analyzed with unmatched one-way ANOVA. P value from multiple comparison test against control are labeled on graph. (E) was analyzed with two-way matched ANOVA, with P value from Sidak’s multiple comparison test labeled on graph. (**F-G**) Asymmetry indices (F) and number of puncta (G) for ASE>AWC iBLINC in indicated conditioning reagents. N = 20 for Naïve, 43 for NaCl, and 24 for glycerol. (F) analyzed with unmatched one-way ANOVA. P value from multiple comparison test against Naïve are labeled on graph. (G) analyzed with two-way matched ANOVA, with P value from Sidak’s multiple comparison test labeled on graph. (**H**) Number of AFD>AIY iBLINC puncta for indicated conditioning. N = 22 (left) and 20 (right) for Naïve, 26 (left) and 27 (right) for 100mM NaCl, 25 for (left) and 26 (right) 0mM NaCl, 23 (left) and 25 (right). Analyzed with two-way unmatched ANOVA, with p value from Sidak’s multiple comparison test labeled on graph.

**Fig. S2.**
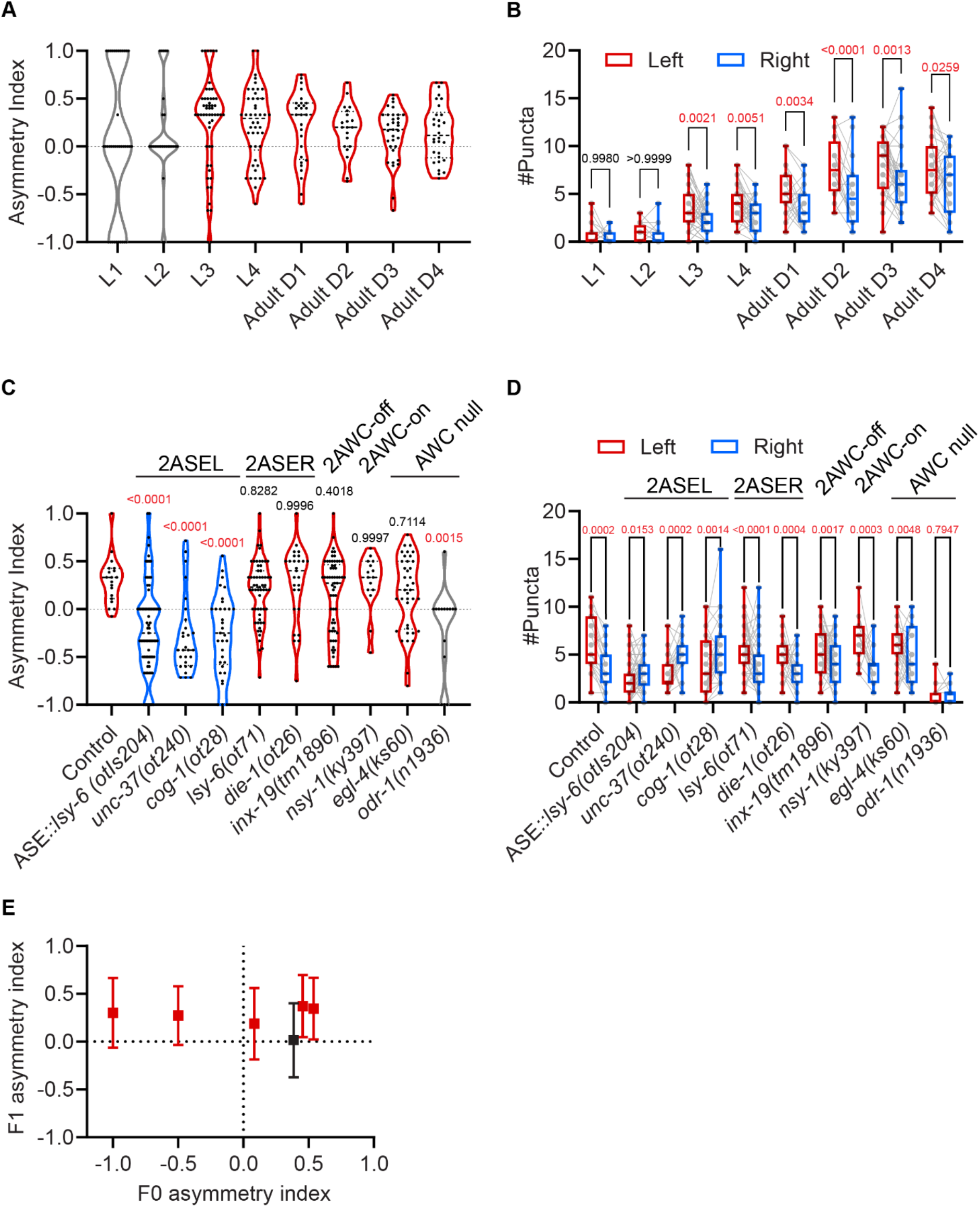
(**A-B**) Asymmetry indices (A) and number of puncta (B) for ASE>AWC iBLINC at indicated ages. N = 38 for L1, 27 for L2, 54 for L3, 46 for L4, 28 for 1 day adult (Adult D1), 20 for 2 day adult (Adult D2), 33 for 3 day adult (Adult D3), 30 for 4 day adult (Adult D4). (B) analyzed with two-way matched ANOVA, with p value from Sidak’s multiple comparison test labeled on graph. (**C-D**) Asymmetry indices (C) and number of puncta (D) for ASE>AWC iBLINC with indicated genotype backgrounds. N = 19 for control, 75 for ASE::lsy-6, 28 for unc-37, 33 for cog-1, 59 for lsy-6, 27 for dis-1, 62 for inx-19, 15 for nsy-1, 38 for egl-4, and 15 for odr-1. (C) analyzed with unmatched one-way ANOVA. P value from multiple comparison test against control are labeled on graph. (D) was analyzed with two-way matched ANOVA, with p value from Sidak’s multiple comparison test labeled on graph. (**E**) Asymmetry indices of F1 animals plotted against the asymmetry indices of their individual maternal F0 animals. Error bars indicate standard deviation. Red indicates significantly left-biased, black indicates no bias. 2-way matched ANOVA with Sidak’s multiple comparison test comparing left and right. For more detail statistic report please refer to Table S10

**Fig. S3.**
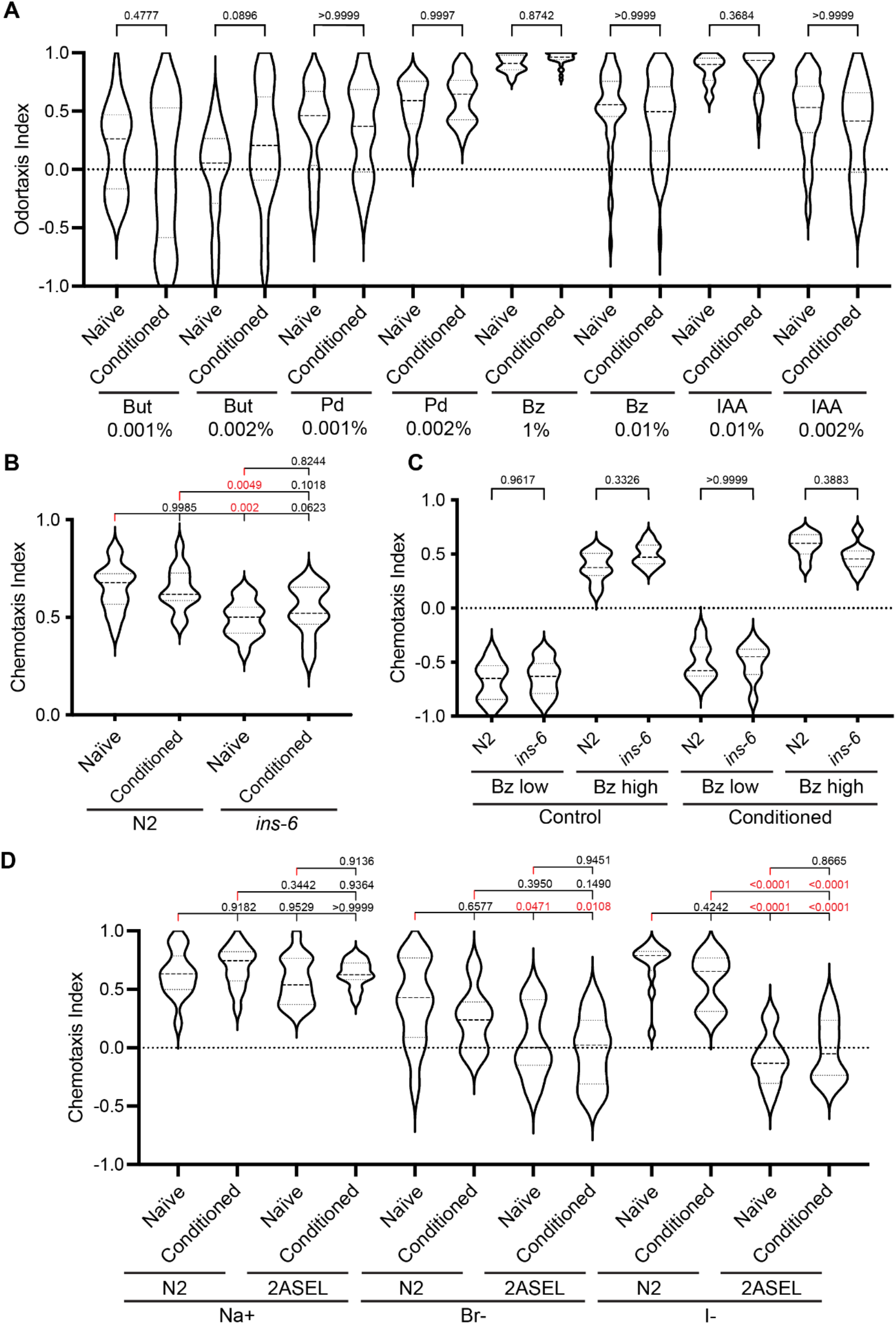
**(A)** Odortaxis indices comparing Control to Conditioned animals’ (12 hrs 100 mM NaCl) attraction to AWC sensed odorants spotted at the indicated concentrations. But: butanone, Pen: pentanedione, Bz: Benzaldehyde, IAA: isoamyl alcohol. Analyzed with unmatched one-way ANOVA. P value from Tukey’s multiple comparisons test labeled on graph. **(B)** Chemotaxis indices indicating N2 and *ins-6(tm2416)* animals’ attraction to a 750 mM NaCl gradient. Conditioning of animals was for 12 hours on plates with 100 mM NaCl. Analyzed with unmatched one-way ANOVA. P value from Tukey’s multiple comparisons test labeled on graph. **(C)** Chemotaxis indices indicating N2 and *ins-6(tm2416)* animals’ attraction to a 100 mM NaCl gradient, with Bz (benzaldehyde) spotted either at the peak of the NaCl gradient (Bz high) or at the opposite point (Bz low). Conditioning of animals was for 12 hours on plates with 100 mM NaCl. Analyzed with unmatched one-way ANOVA. P value from Tukey’s multiple comparisons test labeled on graph **(D)** Chemotaxis indices indicating N2 and 2ASEL mutant (*cog-1(ot28)*) animals’ attraction to ASER-sensed ions, Na+ sodium, Br-bromide, and I-iodide. Analyzed with unmatched one-way ANOVA for each testing reagent. P value from Tukey’s multiple comparisons test labeled on graph.

**Fig. S4.**
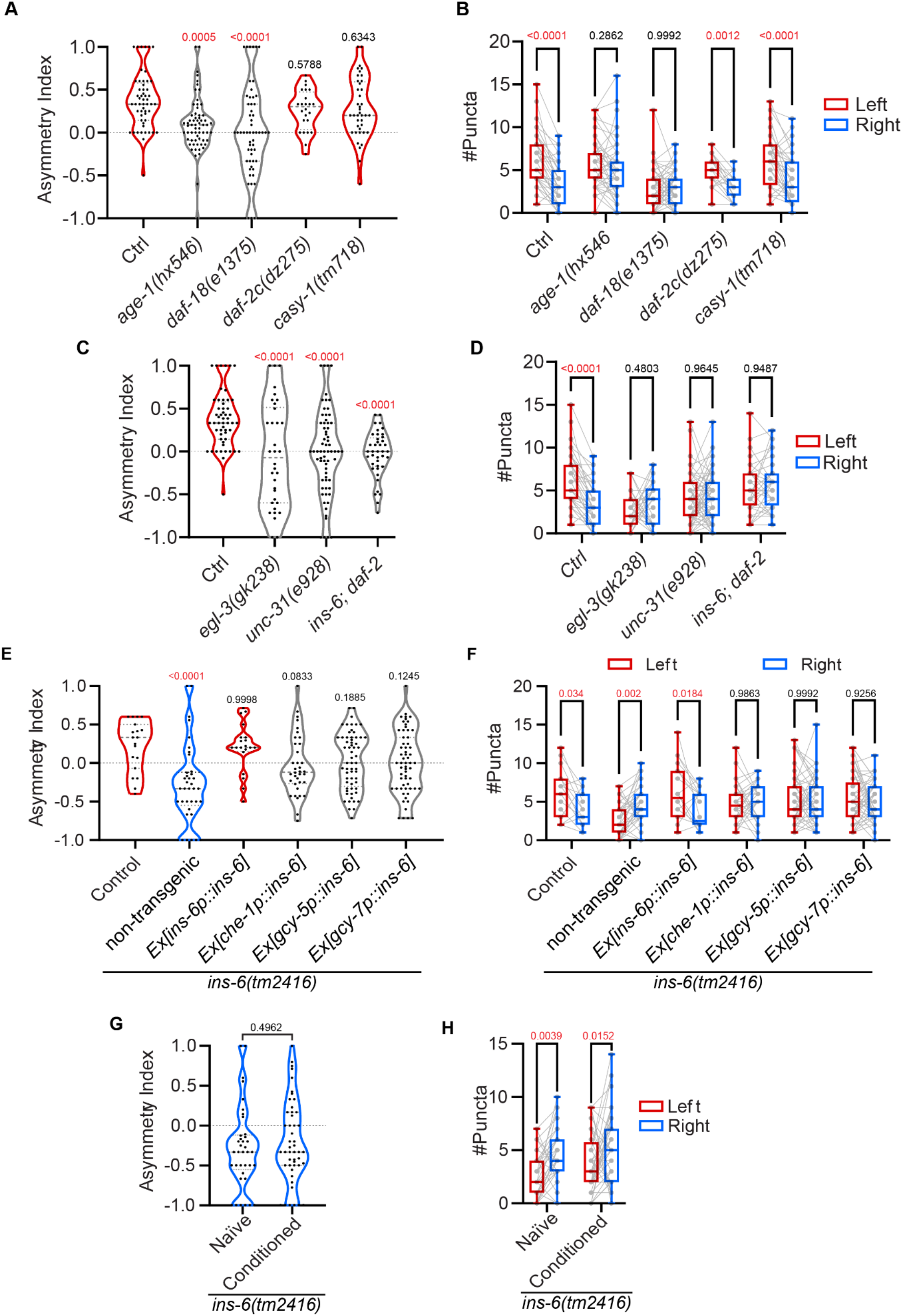
(**A-B**) Asymmetry indices (A) and number of puncta (B) for ASE>AWC iBLINC with indicated genotype backgrounds. N = 53 for Ctrl, 63 for *age-1(hx546)*, 58 for *daf-18(e1375)*, 24 for *daf-2c(dz275)*, 40 for *casy-1(tm718)*. (A) analyzed with unmatched one-way ANOVA. P value from multiple comparison test against control are labeled on graph. (B) analyzed with two-way matched ANOVA, with p value from Sidak’s multiple comparison test labeled on graph. (**C-D**) Asymmetry indices (C) and number of puncta (D) for ASE>AWC iBLINC with indicated genotype backgrounds. N = 53 for Ctrl, 34 for *egl-3(gk238)*, 67 for *unc-31(e928)*, 40 for *ins-6; daf-2*. (C) analyzed with unmatched one-way ANOVA. P value from multiple comparison test against control are labeled on graph. (D) analyzed with two-way matched ANOVA, with P value from Sidak’s multiple comparison test labeled on graph. (**E-F**) Asymmetry indices (E) and number of puncta (F) for ASE>AWC iBLINC in indicated rescue background. N = 19 for Control, 37 for *ins-6 (tm2416)*, 20 for *ins-6p::ins-6*, 36 for *che-1p::ins-6*, 61 for *gcy-5p::ins-6*, and 61 for *gcy-7p::ins-6*. (E) analyzed with unmatched one-way ANOVA. P value from multiple comparison test against control are labeled on graph. (F) was analyzed with two-way matched ANOVA, with p value from Sidak’s multiple comparison test labeled on graph. (**G-H**) Asymmetry indices (G) and number of puncta (H) for ASE>AWC iBLINC in *ins-6(tm2416)* mutant with or without 100mM NaCl conditioning for 12 hours. N = 37 for Naïve, 48 for Conditioned. (G) was analyzed with unmatched two-tailed t-test. (H) was analyzed with two-way matched ANOVA, with p value from Sidak’s multiple comparison test labeled on graph.

**Fig. S5.**
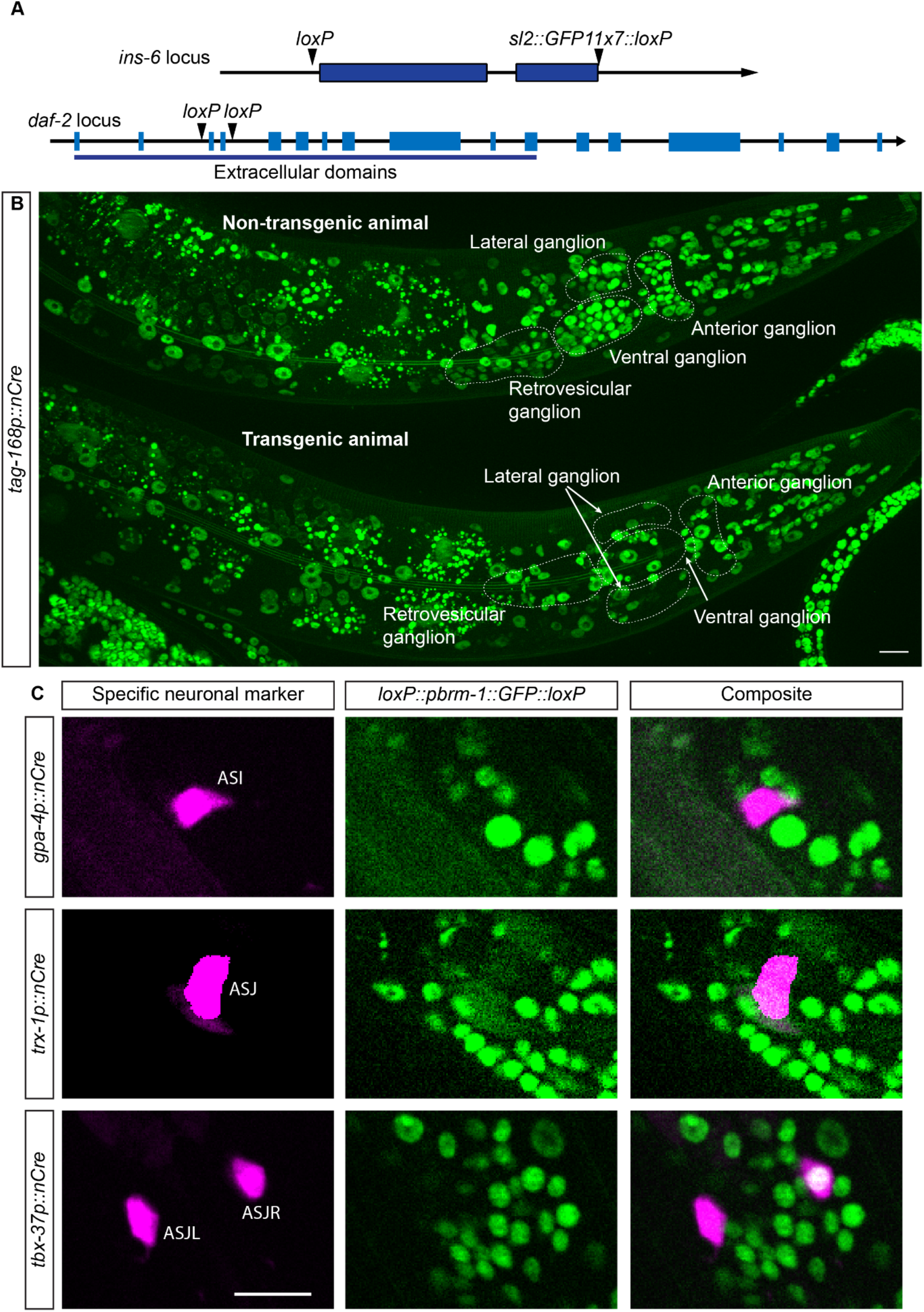
**(A)** Schematic illustrating the location of *LoxP* site inserted to the *ins-6* and *daf-2* locus **(B)** Image of *pbrm-1(rd31[loxP+pbrm-1::GFP+loxP])+dzEx[tag-168p::nCre]*, showing successful Cre-Lox recombination in neuronal cells **(C)** Images of *pbrm-1(rd31[loxP+pbrm-1::GFP+loxP])* with indicated neuronal specific expression of *nCre*, showing successful specific Cre-Lox recombination in the indicated cells

**Fig. S6.**
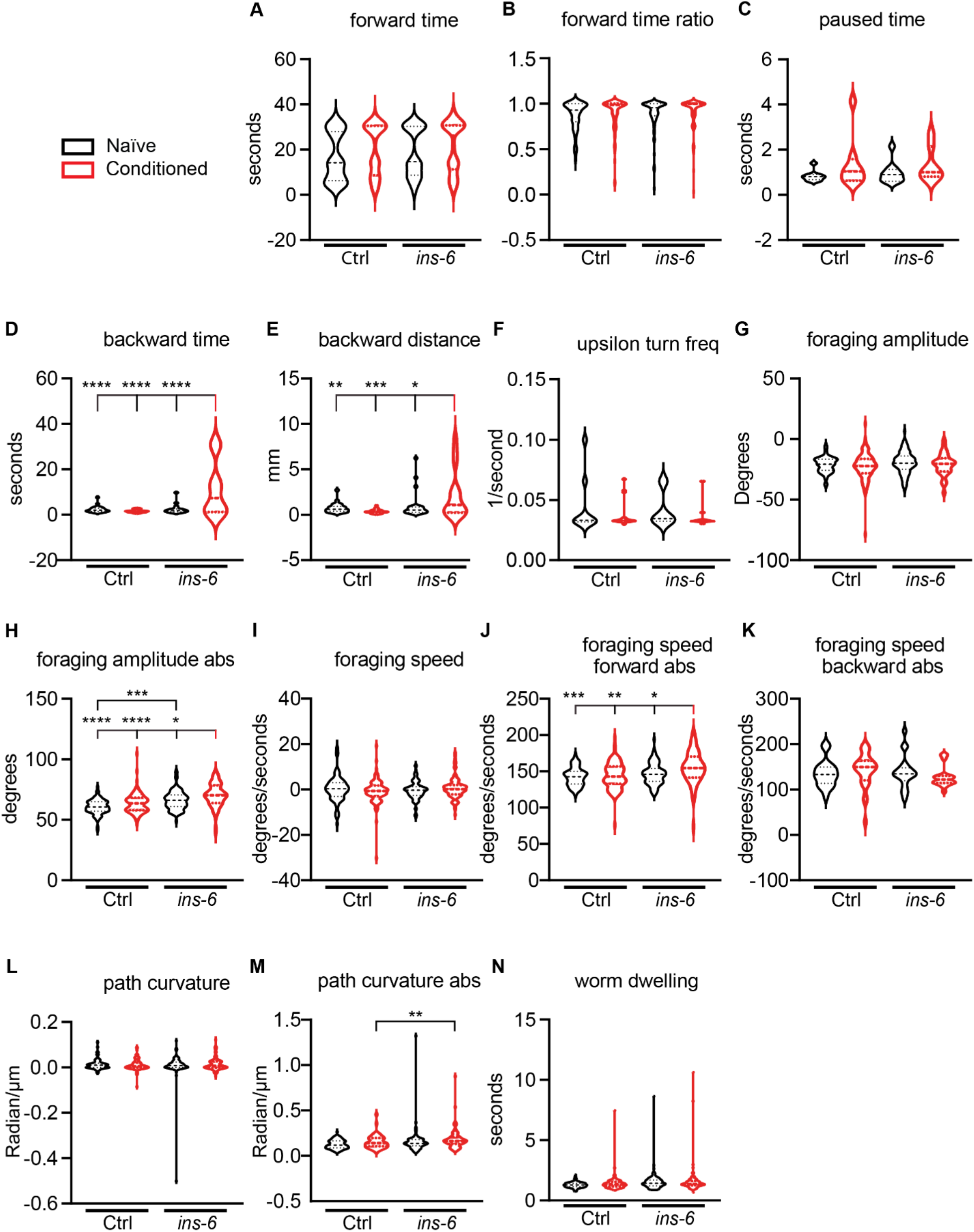
Behavioral parameter extracted from single worm tracking through TierPsy Multi-Worm tracker for naïve and conditioned animals in wildtype or ins-6(tm2416) background. N = 76 for naïve control, 81 for control conditioned, 99 for control *ins-6* mutant, and 81 for conditioned *ins-6* mutant. The details of each behavioral feature are described here. https://github.com/ver228/tierpsy-tracker/blob/master/docs/OUTPUTS.md. Analyzed with one-way ANOVA. For detailed statistic report, see Table S10 in Extended Data

### Supplementary Tables

**Table S1.** The postsynaptic partners, number of synapses and asymmetric indices of ASEL and ASER.

| Animal | N2U |  | JSH |  | n930 |  |
| --- | --- | --- | --- | --- | --- | --- |
|  | ASEL | ASER | ASEL | ASER | ASEL | ASER |
| AFD | 3 | 5 | 1 | 3 | 0 | 1 |
| AIA | 4 | 9 | 4 | 3 | 0 | 1 |
| AIB | 19 | 20 | 11 | 5 | 3 | 4 |
| AIY | 32 | 28 | 8 | 12 | 4 | 11 |
| ASH | 0 | 2 | 1 | 0 | 0 | 0 |
| AWC | 16 | 2 | 4 | 6 | 6 | 0 |

Data extracted from Cook et al (2019) on wormwiring.com, showing only chemical postsynaptic partners that have more than 1 synapses scored from either side.

**Table S2.** List of strains use.

| Strain | Genotype |
| --- | --- |
| N2 |  |
| CB1370 | *daf-2(e1370)* |
| CB1375 | *daf-18(e1375) IV* |
| DA509 | *unc-31(e928) IV* |
| FK163 | *cam-1(ks52) II* |
| FK223 | *egl-4(ks60) IV* |
| FX02416 | *ins-6(tm2416)* |
| HT1593 | *unc-119(ed3) III* |
| OH16024 | *daf-16(ot971[daf-16::GFP]) I* |
| OH2225 | *unc-4(e120) die-1(ot26) II* |
| OH2535 | *lsy-6(ot71) V* |
| OH3903 | *otIs114 I; cog-1(ot28)* |
| OH7805 | *otIs204* |
| OH9305 | *unc-37(ot240) otIs114 I* |
| PR672 | *che-1(p672) I* |
| RA661 | *pbrm-1(rd31[loxP+pbrm-1::GFP+loxP]) I* |
| TJ1052 | *age-1(hx546)* |
| VC461 | *egl-3(gk238) V* |
| EB2443 | *dzIs68 X [AFD>AIY iBLINC]* |
| EB2806 | *dzIs86 I [ASE>AWC iBLINC]; ntlIs1 him-5(e1490) V* |
| EB2835 | *dzIs86 I; otIs204* |
| EB2877 | *dzIs86 I; ins-6(tm2416) II; ntlIs1 V* |
| EB2878 | *dzIs86 I; ntlIs1 lsy-6 (ot71) V* |
| EB2955 | *dzEx1561; dzIs86 I; ntlIs1 him-5(e1490) V* |
| EB2977 | *inx-19(tm1896) dzIs86 I; ntlIs1 him-5(e1490) V* |
| EB3207 | *dzEx1709; dzIs86 I; ins-6(tm2416) II; ntlIs1 V* |
| EB3215 | *dzEx1717; dzIs86 I; ins-6(tm2416) II; ntlIs1 V* |
| EB3220 | *dzEx1722; dzIs86 I; ins-6(tm2416) II; ntlIs1 V* |
| EB3488 | *unc-119() I; dzEx1847* |
| EB3602 | *dzIs86 I; unc-37(ot240) otIs114 I; ntlIs1 him-5(e1490) V* |
| EB3603 | *dzIs86 I; ntlIs1 him-5(e1490) V; egl-4(ks60) IV* |
| EB3604 | *dzIs86 I; cam-1 (ks52) II; ntlIs1 him-5(e1490) V* |
| EB3605 | *dzIs86 I; che-1 (p672) I; ntlIs1 him-5(e1490) V* |
| EB3606 | *dzIs86 I; cog-1(ot28) II* |
| EB3607 | *dzIs86 I; egl-3(gk238) V* |
| EB3608 | *dzIs86 I; unc-4(e120) die-1(ot26) II; ntlIs1 him-5(e1490) V* |
| EB3631 | *dzIs86 I; che-1 (p674) I; ntlIs1 him-5(e1490) V* |
| EB3916 | *dzIs86 I; daf-2(dz275) III* |
| EB3950 | *ins-6(dz280[loxP::ins-6::sl2::nls::GFP11x7::loxP]) II* |
| EB3977 | *dzIs86 I; ins-6(dz280) II* |
| EB3979 | *dzIs86 I; ins-6(dz280) II; dzEx2036* |
| EB3980 | *dzIs86 I; ins-6(dz280) II; dzEx2029* |
| EB3981 | *dzIs86 I; ins-6(dz280) II; dzEx2050* |
| EB4025 | *dzIs86 I; ins-6(dz280) II; dzEx2050* |
| EB4050 | *dzIs86 I; ins-6(dz280) II; dzEx2064* |
| EB4077 | *dzIs86 I; ins-6(tm2416) II; ins-22(ok3616) III* |
| EB4083 | *dzIs86 I; daf-2(dz286) III; dzEx2078* |
| EB4084 | *dzIs86 I; daf-2(dz286) III; dzEx2082* |
| EB4085 | *dzIs86 I; daf-2(dz286) III; dzEx2085* |
| EB4086 | *dzIs86 I; daf-2(dz286)* |
| EB4087 | *dzIs86 I; unc-31(e928) IV* |
| EB4313 | *dzIs86 I; age-1(hx546)* |
| EB4318 | *dzIs86 I; ins-6(tm2614) II; daf-2(e1370) III* |
| EB4353 | *dzIs86 I; daf-2(e1370) III* |
| EB4373 | *daf-16(ot971[daf-16::GFP]) I; dzEx2181* |
| EB4387 | *dzIs86 I; dzEx2184* |
| EB4397 | *dzEx2188* |
| EB4434 | *dzIs127 [flp-6p4::GFP::cla-1+flp-6p4::tagBFP]* |
| EB4440 | *pbrm-1(rd31[loxP+pbrm-1::GFP+loxP]) I; dzEx2195* |
| EB4443 | *pbrm-1(rd31[loxP+pbrm-1::GFP+loxP]) I; dzEx2198* |
| EB4448 | *dzIs86 I; daf-18(e1375) IV* |
| EB4449 | *pbrm-1(rd31[loxP+pbrm-1::GFP+loxP]) I; dzEx2201* |
| EB4455 | *dzIs86 I; dzEx2206* |
| EB4460 | *ins-6(tm2614) II; dzIs127* |
| EB4461 | *dzIs86 I; daf-2(dz286) III; dzEx2211* |
| EB4463 | *dzIs86 I; casy-1(tm718) II* |
| EB4470 | *pbrm-1(rd31[loxP+pbrm-1::GFP+loxP]) I; dzEx2029* |
| EB4472 | *dzIs86 I; dzEx2219* |
| EB4473 | *dzIs86 I; dzEx2220* |
| EB4474 | *dzIs86 I; dzEx2221* |

**Table S3.**
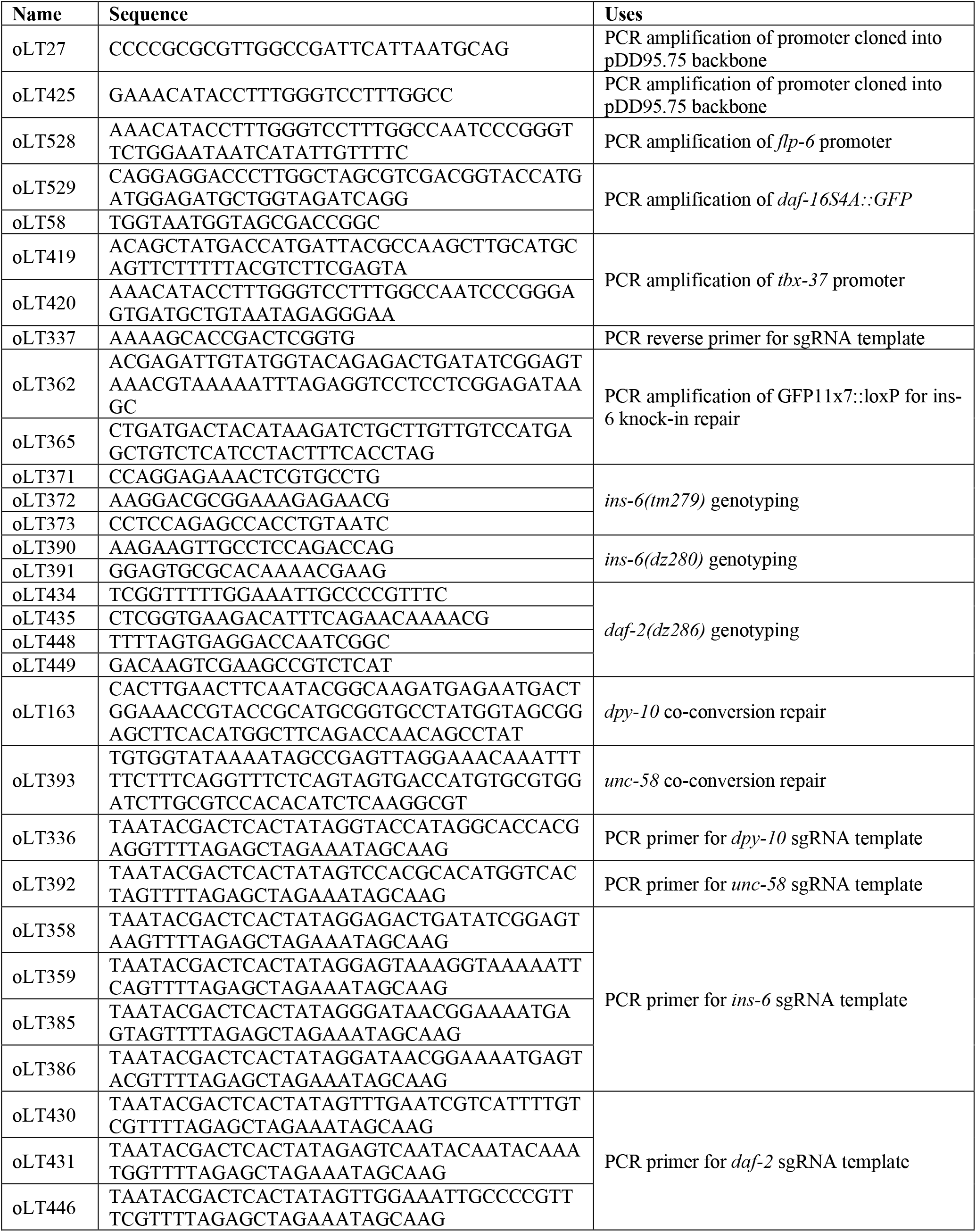
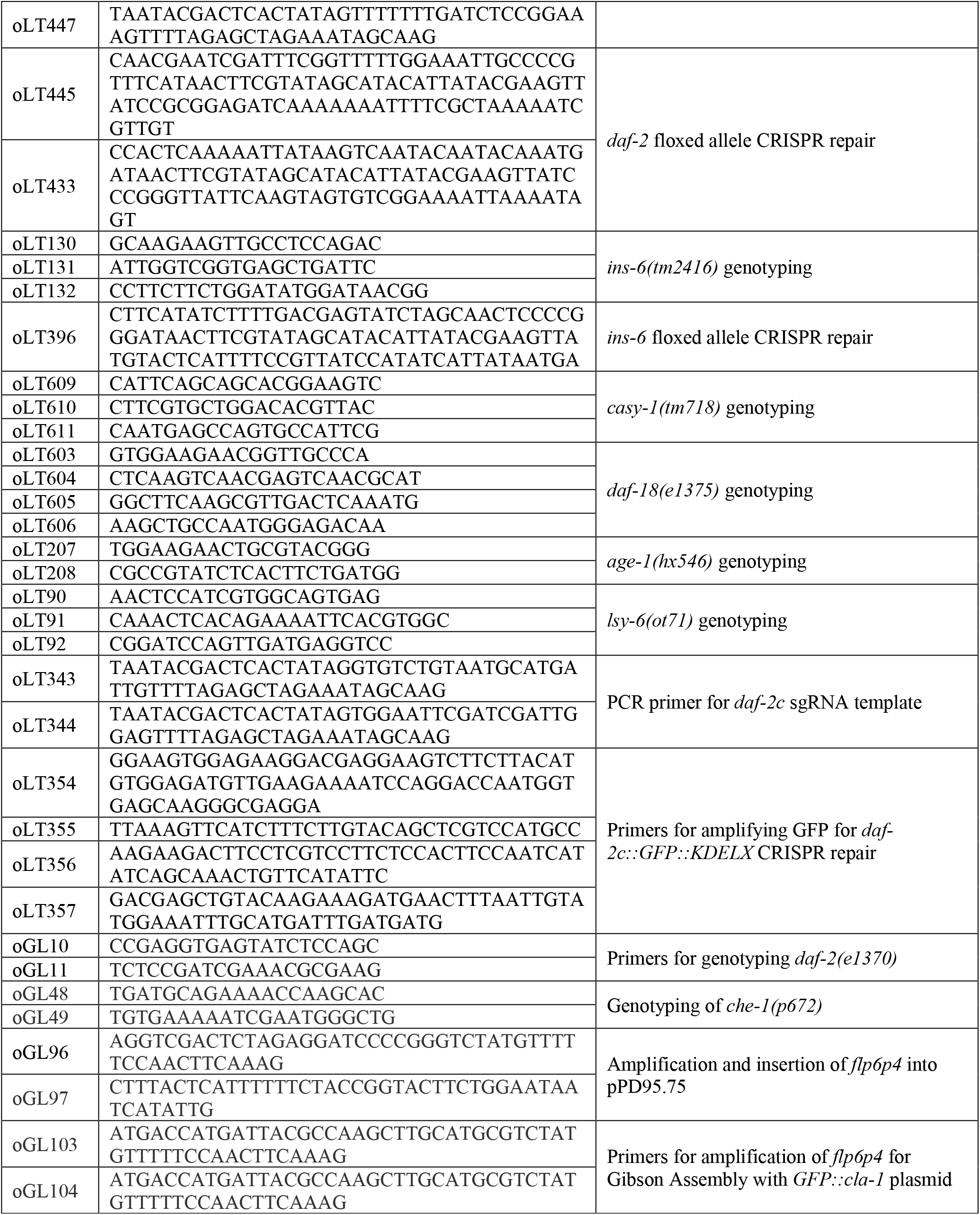
List of oligos used.

**Table S4.** Plasmids generated in previous papers.

| Name | Use |
| --- | --- |
| *pmyo-3p::tagBFP* | Blue muscle injection marker |
| *pDD162* | PCR template for sgRNA transcription template (kind gift from Goldstein Lab) |
| *PK085 (punc-129::3xnovoGFP::cla-1)* | Plasmid used to generate ASE specific CLA-1 active zone marker (kind gift from Dr. Peri Kurshan) |
| *pRF4* | Roller injection marker |
| *pmyo-3p::GFP* | green muscle injection marker |
| *pelt-2p::elt-2::GFP* | green intestine nucleus injection marker |
| *pSF11(ptag-168p::nCre)* | pan-neuronal cell-specific knockouts (kind gift from Dr Dennis Kim) |
| *pZH59 (ptrx-1p::nCre)* | ASJ cell-specific knockouts (kind gift from Dr Dennis Kim) |
| *ges-1p::daf-16S4A::gfp* | Overactive *daf-16S4A::gfp* template (kind gift from Dr Oliver Hobert) |
| *ins-6 [30042]::S0001_pR6K_Amp_2xTY1ce_EGFP_FRT_rpsl_neo_FRT_3xFlag* | Translational reporter for INS-6 (Source Bioscience) |

**Table S5.** Plasmids generated for this paper.

| Name | Use | Construction Method |
| --- | --- | --- |
| *flp-6p4::3xnovoGFP::cla-1* | ASE pre-synaptic marker | *flp-6* promoter was amplified from genomic DNA with the primers oGL103 and oGL104. The resulting PCR product was inserted into PK085 digested with SphI/BssHI through Gibson Assembly. |
| *flp-6p4::tagBFP* | ASE blue marker | *flp-6* promoter was amplified from *pflp-6::gfp* with primers oLT27 and oLT528. The resulting PCR product was inserted into PvuII/XmaI-digested *pmyo-3p::tagBFP* through Gibson Assembly |
| *rab-3p::daf-16S4A::gfp* | pan-neuronal expression of overactive daf-16 | *daf-16S4A::GFP* was amplified from *ges-1p::daf-16S4A::gfp* with primers oLT529 and oLT58. The resulting PCR product was inserted into KpnI/EcoRI-digested *prab-3p::gfp* through Gibson Assembly |
| *gpa-4p::nCre* | ASI cell-specific knockouts | *gpa-4* promoter was amplified from *pgpa-4::tagBFP* with primers oLT27 and oLT425. The resulting PCR product was inserted into PvuII/XmaI-digested *ptrx-1p::nCre* through Gibson Assembly` |
| *tbx-37p::nCre* | ASJL lineage cell-specific knockouts | *tbx-37* promoter was amplified from genomic DNA with primers oLT419 and oLT420. The resulting PCR product was inserted into PvuII/XmaI-digested *ptrx-1p::nCre* through Gibson Assembly` |
| *pflp-6p4::nCre* | ASE cell-specific knockouts | *flp-6* promoter was amplified from *pflp-6::gfp* with primers oLT27 and oLT528. The resulting PCR product was inserted into PvuII/XmaI-digested *ptrx-1p::nCre* through Gibson Assembly` |
| *ceh-36p2d1::nCre* | AWC cell-specific knockouts | *ceh-36p2d1* promoter was amplified from *pceh-36p2d1::tagBFP* with primers oLT27 and oLT425. The resulting PCR product was inserted into PvuII/XmaI-digested *trx-1p::nCre* through Gibson Assembly` |
| *gcy-5p::nCre* | ASER cell-specific knockouts | *gcy-5* promoter was amplified from *gcy-5p::gfp* with primers oLT27 and oLT425. The resulting PCR product was inserted into PvuII/XmaI-digested *trx-1p::nCre* through Gibson Assembly` |
| *gcy-7p::nCre* | ASEL cell-specific knockouts | *gcy-7* promoter was amplified from *gcy-7p::gfp* with primers oLT27 and oLT425. The resulting PCR product was inserted into PvuII/XmaI-digested *ptrx-1p::nCre* through Gibson Assembly` |
| *flp-6p4::daf-16S4A::gfp* | expression of overactive daf-16 in ASE | *daf-16S4A::GFP* was amplified from *ges-1p::daf-16S4A::gfp* with primers oLT529 and oLT58. The resulting PCR product was inserted into KpnI/EcoRI-digested *flp-6p::nls::mCherry* through Gibson Assembly |
| *ceh-36p2d1::daf-16S4A::gfp* | expression of overactive daf-16 in AWC | *daf-16S4A::GFP* was amplified from *ges-1p::daf-16S4A::gfp* with primers oLT529 and oLT58. The resulting PCR product was inserted into KpnI/EcoRI-digested *pceh-36pddl::AP::nlg-1* through Gibson Assembly |
| *flp-6p4::nls::mCherry* | ASE bilateral red nuclear marker | *flp-6* promoter was amplified from *pflp-6::gfp* with primers oLT27 and oLT528. The resulting PCR product was inserted into PvuII/XmaI-digested *act-4p::nls::mCherry* through Gibson Assembly` |
| *flp-6p4::gfp* | ASE bilateral green marker | *flp-6* promoter was amplified from genomic DNA with the primers oGL96 and oGL97. The resulting PCR product was inserted into pPD95.75 digested with XmaI and AgeI using Gibson Assembly. |

**Table S6.** List of Extrachromosomal array generated.

| Strain | Array | Use | Plasmids contained | Line number |
| --- | --- | --- | --- | --- |
| EB4472 | *dzEx2219* | pan-neuronal expression of overactive daf-16 | *25 ng/μl rab-3p::daf-16S4A::GFP; 10 ng/μl myo3p::BFP* | 1 |
| EB4473 | *dzEx2220* |  |  | 2 |
| EB4474 | *dzEx2221* |  |  | 3 |
| EB3488 | *dzEx1847* | *ins-6* translational reporter | *5 ng/μl (ins-6[30042]::S0001_pR6K_Amp_2xTY1ce_EGFP_FRT_rpsl neo FRT 3xFlag)dFRT::unc-119-Nat* | 1 |
| EB3958 | *dzEx2036* | ASJ specific floxed allele knockout | *25 ng/μl trx-1p::nCre, 5 ng/μl myo-3p::tagBFP* | 1 |
| EB3959 | *dzEx2037* |  |  | 2 |
| EB3960 | *dzEx2038* |  |  | 3 |
| EB3961 | *dzEx2039* |  |  | 4 |
| EB3945 | *dzEx2029* | pan-neuronal floxed allele knockout | *5 ng/μl myo-3p::tagBFP, 20 ng/μl tag-168p::nCre* | 1 |
| EB3946 | *dzEx2030* |  |  | 2 |
| EB3947 | *dzEx2029* |  |  | 3 |
| EB3948 | *dzEx2031* |  |  | 4 |
| EB3949 | *dzEx2029* |  |  | 5 |
| EB3981 | *dzEx2050* | ASI specific flox allele knockout | *25 ng/μl gpa-4p::nCre, 5 ng/μl myo-3p::GFP* | 1 |
| EB3982 | *dzEx2051* |  |  | 2 |
| EB3983 | *dzEx2052* |  |  | 3 |
| EB4025 | *dzEx2064* | ASI+ASJ specific flox allele knockout | *40 ng/μl trx-1p::nCre, 40 ng/μl gpa-4p::nCre, 10 ng/μl elt-2p::GFP* | 1 |
| EB4026 | *dzEx2065* |  |  | 2 |
| EB4027 | *dzEx2066* |  |  | 3 |
| EB4050 | *dzEx2075* | ABa lineage flox allele knockout | *50 ng/μl tbx-37p::nCre, 5 ng/μl myo-3p::GFP* | 1 |
| EB4051 | *dzEx2076* |  |  | 2 |
| EB4049 | *dzEx2074* |  |  | 3 |
| EB4052 | *dzEx2077* |  |  | 4 |
| EB4060 | *dzEx2078* | ASI specific flox allele knockout | *5 ng/μl myo-3p::gfp, 25 ng/μl gcy-5p::nCre* | 1 |
| EB4061 | *dzEx2079* |  |  | 2 |
| EB4062 | *dzEx2080* |  |  | 3 |
| EB4067 | *dzEx2085* | ASI specific flox allele knockout | *5 ng/μl myo-3p::gfp, 25 ng/μl ceh-36p2del1::nCre* | 1 |
| EB4068 | *dzEx2086* |  |  | 2 |
| EB4069 | *dzEx2087* |  |  | 3 |
| EB4461 | *dzEx2211* | ASI specific flox allele knockout | *25 ng/μl flp-6::nCre; 10 ng/μl myo3p::BFP* | 1 |
| EB4462 | *dzEx2212* |  |  | 2 |
| EB4375 | *dzEx2183* | ASI specific flox allele knockout | *25 ng/μl flp-6::nls::mCherry, 25 ng/μl flp-6::tagBFP, 40 ng/μl pRF4* | 1 |
| EB4373 | *dzEx2181* |  |  | 2 |
| EB4374 | *dzEx2182* |  |  | 3 |
| EB4455 | *dzEx2206* |  | *25 ng/μl flp-6p::daf-16S4A::GFP; 10 ng/μl myo-3p::BFP* | 1 |
| EB4456 | *dzEx2209* |  |  | 2 |
| EB4457 | *dzEx2210* | ASE expression of overactive *daf-16* |  | 3 |
| EB4387 | *dzEx2184* | AWC expression of overactive *daf-16* | *25ng/μl ceh-36p2d1::daf-16S4A::GFP, 10 ng/μl myo-3p::tagBFP* | 1 |
| EB3207 | *dzEx1709* | *ins-6* rescue from ASER | *5ng/μl gcy-5p::ins-6 (PCR product) + 10 ng/μl elt-2p::GFP* | 1 |
| EB3208 | *dzEx1710* |  |  | 2 |
| EB3209 | *dzEx1711* |  |  | 3 |
| EB3210 | *dzEx1712* |  |  | 4 |
| EB3211 | *dzEx1713* |  |  | 5 |
| EB3212 | *dzEx1714* |  |  | 6 |
| EB3213 | *dzEx1715* |  |  | 7 |
| EB3214 | *dzEx1716* |  |  | 8 |
| EB3215 | *dzEx1717* | *ins-6* rescue from ASEL | *5ng/μl gcy-7p::ins-6 (PCR product) + 10 ng/μl elt-2p::GFP* | 1 |
| EB3216 | *dzEx1718* |  |  | 2 |
| EB3217 | *dzEx1719* |  |  | 3 |
| EB3218 | *dzEx1720* |  |  | 4 |
| EB3219 | *dzEx1721* |  |  | 5 |
| EB3220 | *dzEx1722* | *ins-6* rescue from ASE | *20 ng/μl che-1p::ins-6 (PCR product) + 10 ng/μl elt-2p::GFP* | 1 |
| EB3221 | *dzEx1723* |  |  | 2 |
| EB3222 | *dzEx1724* |  |  | 3 |
| EB3223 | *dzEx1725* |  |  | 4 |
| EB3224 | *dzEx1726* |  |  | 5 |
| EB3225 | *dzEx1727* |  |  | 6 |
| EB4440 | *dzEx2195* | ASI specific flox allele knockout | *25 ng/μl gpa-4p::nCre, 10 ng/μl gpa-4p::tagBFP 10 ng/μl myo-3p::BFP* | 1 |
| EB4441 | *dzEx2196* |  |  | 2 |
| EB4442 | *dzEx2197* |  |  | 3 |
| EB4443 | *dzEx2198* | ASJ specific flox allele knockout | *25 ng/μl trx-1p::nCre, 10 ng/μl trx-1p::tagBFP, 10 ng/μl myo-3p::BFP* | 1 |
| EB4444 | *dzEx2199* |  |  | 2 |
| EB4445 | *dzEx2200* |  |  | 3 |
| EB4449 | *dzEx2201* | ABa lineage flox allele knockout | *25 ng/μl tbx-37p::nCre, 10 ng/μl trx-1p::tagBFP, 10 ng/μl myo-3p::BFP* | 1 |
| EB4064 | *dzEx2082* | ASI specific flox allele knockout | *5 ng/μl myo-3prom::tagBFP, 25 ng/μl gcy-7prom:nCre* | 1 |
| EB4065 | *dzEx2083* |  |  | 2 |
| EB4066 | *dzEx2084* |  |  | 3 |
| EB4397 | *dzEx2188* | *flp-6p::GFP* | *25 ng/μl flp-6prom2::3xnovoGFP::cla-1; 25 ng/μl flp-6prom2::tagBFP* | 1 |

**Table S7.**
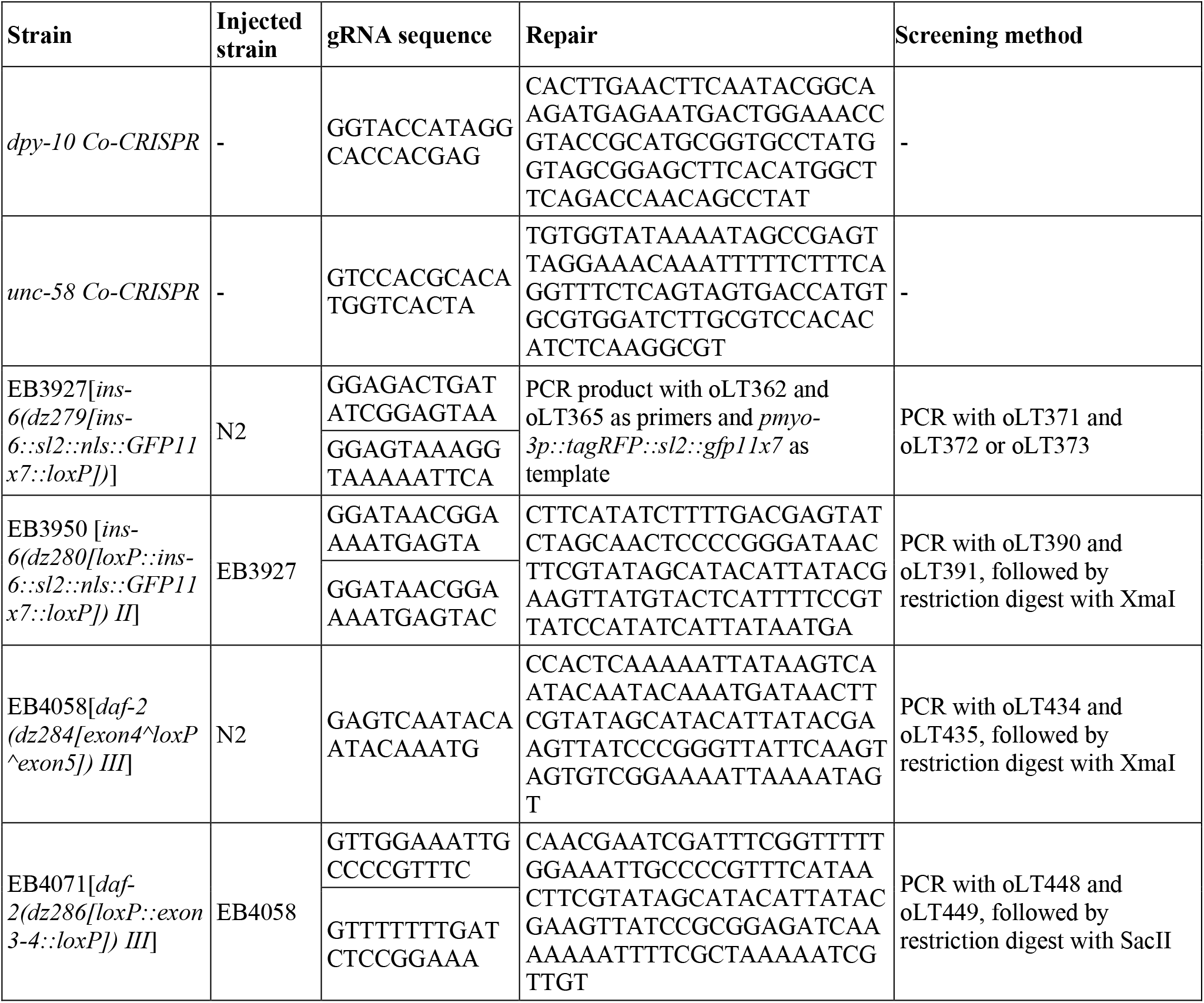
CRISPR strains generated.

**Table S8.** Statistics summary of two-way ANOVA for left-right asymmetry.

| Figure | F (DFn, DFd) |  |  |  | P values |  |  |  |
| --- | --- | --- | --- | --- | --- | --- | --- | --- |
|  | Left vs Right | Experimental Variable | Interaction | Individual animals | Left vs Right | Experimental Variable | Interaction | Individual animals |
| 1H | F (1, 73) = 1.260 | F (1, 73) = 28.48 | F (1, 73) = 26.21 | F (73, 73) = 1.094 | P=0.2653 | P<0.0001 | P<0.0001 | P=0.3508 |
| 1J | F (1, 134) = 0.04031 | F (3, 134) = 6.616 | F (3, 134) = 7.040 | F (134, 134) = 3.027 | P=0.8412 | P=0.0003 | P=0.0002 | P<0.0001 |
| 2C | F (1, 95) = 4.491 | F (1, 95) = 0.6025 | F (1, 95) = 47.27 | F (95, 95) = 1.471 | P=0.0367 | P=0.4396 | P<0.0001 | P=0.0308 |
| 2G | F (1, 369) = 3.025 | F (6, 369) = 7.753 | F (6, 369) = 11.61 | F (369, 369) = 1.076 | P=0.0828 | P<0.0001 | P<0.0001 | P=0.2403 |
| 2I | F (1, 128) = 5.880 | F (6, 128) = 8.220 | F (6, 128) = 5.184 | F (128, 128) = 0.6853 | P=0.0167 | P<0.0001 | P<0.0001 | P=0.9833 |
| 2J (4 HOURS) | F (1, 148) = 15.64 | F (4, 148) = 3.328 | F (4, 148) = 4.781 | F (148, 148) = 1.622 | P=0.0001 | P=0.0121 | P=0.0012 | P=0.0017 |
| 2J (6 HOURS) | F (1, 128) = 0.4623 | F (4, 128) = 8.366 | F (4, 128) = 3.140 | F (128, 128) = 1.066 | P=0.4978 | P<0.0001 | P=0.0168 | P=0.3597 |
| 2J (12 HOURS) | F (1, 193) = 1.630 | F (4, 193) = 23.29 | F (4, 193) = 7.567 | F (193, 193) = 1.126 | P=0.2032 | P<0.0001 | P<0.0001 | P=0.2046 |
| 3D | F (1, 120) = 11.93 | F (2, 120) = 2.186 | F (2, 120) = 8.346 | F (120, 120) = 1.570 | P=0.0008 | P=0.1169 | P=0.0004 | P=0.0070 |
| 3F | F (1, 242) = 66.54 | F (4, 242) = 8.620 | F (4, 242) = 10.12 | F (242, 242) = 2.237 | P<0.0001 | P<0.0001 | P<0.0001 | P<0.0001 |
| 3H | F (1, 273) = 3.033 | F (5, 273) = 9.621 | F (5, 273) = 12.78 | F (273, 273) = 2.192 | P=0.0827 | P<0.0001 | P<0.0001 | P<0.0001 |
| 4C | F (1, 88) = 2.479 | F (1, 88) = 3.403 | F (1, 88) = 77.82 | F (88, 88) = 2.919 | P=0.1190 | P=0.0685 | P<0.0001 | P<0.0001 |
| 4E | F (1, 29) = 0.1527 | F (1, 29) = 0.07135 | F (1, 29) = 14.13 | F (29, 29) = 1.664 | P=0.6988 | P=0.7913 | P=0.0008 | P=0.0883 |
| 4G | F (1, 146) = 0.0021 | F (4, 146) = 12.63 | F (4, 146) = 20.57 | F (146, 146) = 3.204 | P=0.9634 | P<0.0001 | P<0.0001 | P<0.0001 |
| 4J | F (1, 70) = 0.9611 | F (1, 70) = 6.421 | F (1, 70) = 53.26 | F (70, 70) = 2.856 | P=0.3303 | P=0.0135 | P<0.0001 | P<0.0001 |
| S1C | F (1, 226) = 148.9 | F (11, 226) = 10.64 | F (11, 226) = 4.469 | F (226, 226) = 1.305 | P<0.0001 | P<0.0001 | P<0.0001 | P=0.0229 |
| S1E | F (1, 123) = 3.894 | F (2, 123) = 9.449 | F (2, 123) = 9.559 | F (123, 123) = 0.6866 | P=0.0507 | P=0.0002 | P=0.0001 | P=0.9810 |
| S1G | F (1, 138) = 8.853 | F (4, 138) = 3.141 | F (4, 138) = 10.92 | F (138, 138) = 1.212 | P=0.0035 | P=0.0166 | P<0.0001 | P=0.1296 |
| S2B | F (1, 269) = 65.61 | F (7, 269) = 71.22 | F (7, 269) = 2.583 | F (269, 269) = 1.801 | P<0.0001 | P<0.0001 | P=0.0136 | P<0.0001 |
| S2D | F (1, 364) = 13.86 | F (9, 364) = 17.85 | F (9, 364) = 10.55 | F (364, 364) = 1.325 | P=0.0002 | P<0.0001 | P<0.0001 | P=0.0037 |
| S4B | F (1, 235) = 79.49 | F (4, 235) = 9.551 | F (4, 235) = 11.38 | F (235, 235) = 3.170 | P<0.0001 | P<0.0001 | P<0.0001 | P<0.0001 |
| S4D | F (1, 190) = 3.226 | F (3, 190) = 7.278 | F (3, 190) = 15.04 | F (190, 190) = 2.267 | P=0.0741 | P=0.0001 | P<0.0001 | P<0.0001 |
| S4F | F (1, 228) = 3.104 | F (5, 228) = 2.027 | F (5, 228) = 6.213 | F (228, 228) = 1.837 | P=0.0795 | P=0.0758 | P<0.0001 | P<0.0001 |
| S4H | F (1, 83) = 17.74 | F (1, 83) = 3.180 | F (1, 83) = 0.3630 | F (83, 83) = 1.138 | P<0.0001 | P=0.0782 | P=0.5485 | P=0.2789 |

**Table S9.** Statistic Summary for Figure 2J.

| Time point | Factor analyzed | F(DFn, DFd) | P | Conditioning NaCl Conc. | n | *P* value from Šídák's multiple comparisons test |
| --- | --- | --- | --- | --- | --- | --- |
| 4 hour | Left-right | F (4, 148) = 3.328 | P=0.0121 | 50 mM | 17 | 0.0004 |
|  | Conditioning conc. | F (4, 148) = 4.781 | P=0.0012 | 75 mM | 26 | 0.0049 |
|  | Interaction | F (1, 148) = 15.64 | P=0.0001 | 100 mM | 64 | >0.9999 |
|  | Animal | F (148, 148) = 1.622 | P=0.0017 | 150 mM | 23 | 0.9778 |
|  |  |  |  | 200 mM | 23 | >0.9999 |
| 6 hour | Left-right | F (4, 128) = 3.140 | P=0.0168 | 50 mM | 20 | 0.0432 |
|  | Conditioning conc. | F (4, 128) = 8.366 | P<0.0001 | 75 mM | 16 | 0.9866 |
|  | Interaction | F (1, 128) = 0.4623 | P=0.4978 | 100 mM | 62 | 0.1191 |
|  | Animal | F (128, 128) = 1.066 | P=0.3597 | 150 mM | 16 | 0.9922 |
|  |  |  |  | 200 mM | 19 | >0.9999 |
| 12 hour | Left-right | F (4, 193) = 7.567 | P<0.0001 | 50 mM | 23 | 0.0018 |
|  | Conditioning conc. | F (4, 193) = 23.29 | P<0.0001 | 75 mM | 36 | 0.9969 |
|  | Interaction | F (1, 193) = 1.630 | P=0.2032 | 100 mM | 78 | 0.0258 |
|  | Animal | F (193, 193) = 1.126 | P=0.2046 | 150 mM | 31 | 0.0419 |
|  |  |  |  | 200 mM | 30 | 0.0231 |

**Table S10.** Statistic Summary of behavioral assay for Extended Data Figure 6.

| Parameter | F (DFn, Dfd) | P Value | Tukey's multiple comparison test P values |  |  |  |  |  |
| --- | --- | --- | --- | --- | --- | --- | --- | --- |
|  |  |  | ins6 naive vs. ins6 salt | ins6 naive vs. control naive | ins6 naive vs. control salt | ins6 salt vs. control naive | ins6 salt vs. control salt | control naive vs. control salt |
| Forward Time | F (3, 324) = 2.372 | P=0.0703 | 0.4884 | 0.6114 | 0.8064 | 0.0662 | 0.9525 | 0.1859 |
| Forward time ratio | F (3, 324) = 0.2579 | P=0.8557 | 0.9998 | 0.8727 | 0.9942 | 0.866 | 0.99 | 0.9621 |
| Forward distance | F (3, 333) = 0.9441 | P=0.4195 | 0.6681 | 0.9896 | 0.9679 | 0.5246 | 0.4269 | 0.9991 |
| Paused Time | F (3, 31) = 1.331 | P=0.2820 | 0.6863 | 0.9689 | 0.6194 | 0.4385 | >0.9999 | 0.3692 |
| backward time | F (3, 86) = 14.32 | P<0.0001 | <0.0001 | >0.9999 | 0.9564 | <0.0001 | <0.0001 | 0.9498 |
| backward distance | F (3, 89) = 6.863 | P=0.0003 | 0.0164 | 0.9365 | 0.4778 | 0.0032 | 0.0002 | 0.8274 |
| upsilon turn freq | F (3, 52) = 0.6241 | P=0.6027 | 0.7314 | 0.9945 | 0.88 | 0.6662 | 0.9841 | 0.8116 |
| foraging amplitude | F (3, 337) = 1.488 | P=0.2176 | 0.6477 | 0.6659 | 0.1578 | >0.9999 | 0.8146 | 0.8036 |
| foraging amplitude abs | F (3, 337) = 17.45 | P<0.0001 | 0.0136 | 0.0001 | 0.2768 | <0.0001 | <0.0001 | 0.0773 |
| foraging speed | F (3, 333) = 1.507 | P=0.2124 | 0.4454 | 0.8152 | 0.9498 | 0.9448 | 0.2165 | 0.535 |
| foraging speed forward abs | F (3, 333) = 6.876 | P=0.0002 | 0.0154 | 0.5444 | 0.8066 | 0.0003 | 0.0014 | 0.9735 |
| foraging speed backward abs | F (3, 85) = 0.6530 | P=0.5833 | 0.7688 | 0.9939 | 0.9807 | 0.8735 | 0.5205 | 0.9075 |
| path curvature | F (3, 333) = 0.4655 | P=0.7066 | 0.6932 | 0.8326 | 0.9714 | 0.9963 | 0.9245 | 0.9792 |
| path curvature abs | F (3, 337) = 4.311 | P=0.0053 | 0.2336 | 0.2364 | >0.9999 | 0.0021 | 0.301 | 0.2501 |
| worm dwelling | F (3, 333) = 2.466 | P=0.0622 | 0.9066 | 0.1897 | 0.7997 | 0.0552 | 0.4321 | 0.7156 |

